# NGFR regulates germinal center B-cell activation and negative selection

**DOI:** 10.1101/2023.07.14.548960

**Authors:** Alberto Hernández-Barranco, Marina S. Mazariegos, Vanesa Santos, Eduardo Caleiras, Laura Nogués, Frédéric Mourcin, Simon Léonard, Christelle Oblet, Steve Genebrier, Delphine Rossille, Alberto Benguría, Enrique Vazquez, Ana Dopazo, Alejo Efeyan, Ana Ortega-Molina, Michel Cogne, Karin Tarte, Héctor Peinado

## Abstract

The expression of the Nerve growth factor receptor (NGFR) has been described in follicular dendritic cells (FDCs), the major lymphoid stromal cell (LSC) compartment regulating B-cell activation within germinal centers (GCs). However, the role of NGFR in humoral response is not well defined. In this work, we have studied the effect of *Ngfr* KO in LNs organization and function. *Ngfr* KO led to spontaneous GC formation and expansion of GC B-cell compartment which were related to *Ngfr* depletion in non-hematopoietic radioresistant compartment. In agreement, *Ngfr* KO mice showed alterations in LSC with an increased frequency of FDCs harboring an activated phenotype characterized by the overexpression of CD21/35, MAdCAM-1, and VCAM-1. Moreover, *Ngfr* KO mice showed GC ectopic location, loss of polarization, impaired high-affinity antibody production, and increased circulating autoantibodies. In addition, *Ngfr* KO*/Bcl2* Tg mice displayed increased levels of autoantibodies, higher incidence of autoimmunity, and decreased overall survival. Our work shows that NGFR maintains GC structure and functionality, being involved in the regulation of antibody production and immune tolerance.

## Introduction

The nerve growth factor (NGF) receptor (NGFR), also known as p75^NTR^, belongs to the tumor necrosis factor (TNF) receptor (TNFR) superfamily. NGFR is a low affinity pan-receptor of the neurotrophin family of proteins, which includes NGF,^1^ the brain-derived neurotrophic factor (BDNF),^2^ the neurotrophin-3 (NT3),^3^ and the neurotrophin-4/5 (NT4/5),^4^ as well as their pre-processed immature forms (known as pro-neurotrophins).^5^ Despite being first discovered in the nervous system playing a role in the control of cell survival and apoptosis,^6–8^ NGFR has been later detected in many other organs fulfilling a large variety of functions in different physiological and pathological processes.^9,10^

Out of the many analyzed cell types, high NGFR expression in follicular dendritic cells (FDCs) was reported decades ago.^11,12^ However, its functional role in this context remains poorly explored. FDCs are a subset of lymphoid stromal cells (LSCs)^13–16^ involved in the control of B cell maturation and selection within germinal centers (GCs). In physiological conditions, GCs have a transient nature, playing a critical role in the normal development of humoral immunity in response to pathogens and immunization.^17,18^ GC microstructural organization ensures the correct regulation of the different steps of the GC reaction, with in particular B-cell proliferation and mutations in immunoglobulin variable genes occurring in GC dark zone (DZ), whereas B-cell affinity selection occurs in GC light zone (LZ). This regulation is not only driven by the genetic program of the B-lymphocytes, but also by paracrine signals and cell-to-cell interactions provided by cells in the surrounding microenvironment.^19,20^ In fact, alterations in the microenvironment lead to pathological conditions such as autoimmunity and follicular lymphoma.^20–23^

Here, we have analyzed the role of NGFR in lymph nodes (LNs) and the implications of its loss in the development of autoimmunity. We observed that NGFR expression is modulated upon FDC activation by immunization. We characterized the phenotype of secondary lymphoid organs (SLOs) in a *Ngfr* KO mouse model, finding that these mice had enlarged LNs due to GC hyperplasia and spontaneous activation. Such alterations were reproduced when *Ngfr* was depleted only in the non-hematopoietic compartment, demonstrating a B-cell extrinsic activity. GCs in *Ngfr* KO LNs showed structural and functional abnormalities compared to GCs induced in WT mice upon immunization. These alterations included ectopic GC location, loss of DZ/LZ polarization, and impaired high-affinity antibody production, together with the presence of circulating autoantibodies. FDCs were the main LSC population increased in LNs from naïve *Ngfr* KO mice and their characterization by RNAseq showed an overexpression of FDCs activation markers, including CD21/35, MAdCAM-1 and VCAM-1. We aimed to gain more insights in the regulation of autoimmunity by depleting *Ngfr* in a *Bcl2* transgenic (Tg) mouse model known to decrease the negative selection of B cells by impairing apoptosis.^24^ Double mutant mice had increased levels of autoantibodies in serum concomitant with a higher incidence of severe autoimmune lesions such as lupus-like glomerulonephritis, responsible for the decreased overall survival observed in this model. Overall, our work shows that NGFR regulates B-cell immune responses by preserving peripheric tolerance and preventing autoimmunity.

## Results

### NGFR is modulated in FDCs in response to immunization

We first studied NGFR dynamics in FDCs during an immune response after immunization in WT mice. We injected intra-footpad NP hapten conjugated to Keyhole Limpet Hemocyanin (NP-KLH) to induce GC formation.^25^ We analyzed the expression levels of NGFR *in situ* in CD21/35^+^ FDCs from WT naïve and immunized animals by immunohistofluorescence. Our results revealed that NGFR expression was significantly reduced in FDCs 10 days post-immunization (**Figure 1A, B**), suggesting its functional implication. CD21/35 expression was previously demonstrated to be induced upon activation and polarization of primary follicle FDCs differentiating LZ-FDCs (which exhibit higher CD21/35 levels) from DZ-FDCs. ^26,27^ Thus, we used this marker to differentiate two subpopulations of FDCs in LNs from immunized WT mice by flow cytometry. CD21/35^hi^ FDCs also showed increased levels of other LZ-FDC activation markers such as MAdCAM-1 and VCAM-1 compared to CD21/CD35^lo^ FDCs (**Figure 1 C, D)**. Analysis of NGFR showed that its expression was lower in CD21/35^hi^ FDCs (**Figure 1E**), exhibiting an inverse correlation with the other LZ-FDC activation markers (**Figure 1F**). These data suggested that NGFR modulation is involved in FDC activation and differentiation.

**Figure 1:**
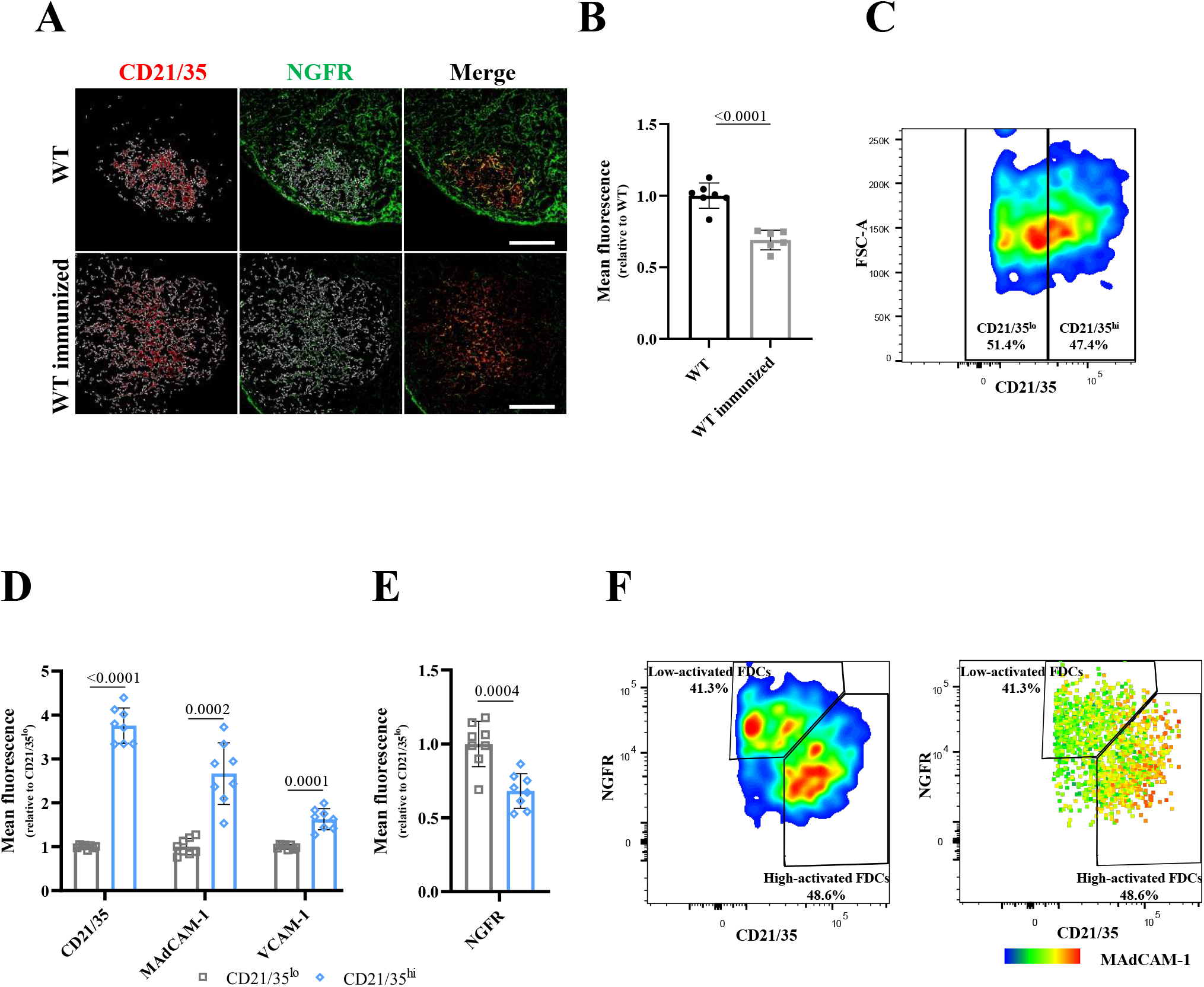
NGFR is modulated in response to immunization. **A**) Representative IF images of CD21/35 (red), and NGFR (green) in GCs from WT and WT immunized PO LNs. White line shows the area selected for quantification based on CD21/35 expression. Scale bar, 100µm. **B**) Quantification by IF of NGFR mean fluorescence in FDCs from WT, and WT immunized PO LNs. Up to 3 individual follicles per LN were analyzed when possible and average values were plotted. n=7 WT and 6 WT immunized LNs from 1 experiment. **C**) Gating strategy used to select CD21/35^hi^ and ^lo^ populations in FDCs from PO LNs 10 days after immunization in WT mice. **D**) Quantification by flow cytometry of the mean fluorescence of the indicated FDCs activation markers in the populations gated in **C.** n=8 from two independent experiments**. E**) Quantification by flow cytometry of NGFR mean fluorescence in the populations gated in **C**. **F**) Representative pseudocolor plot (left panel) and heatmap statistics plot (right panel) of immunized FDCs. The heatmap shows MaDCAM-1 mean fluorescence. Representative plots in **C** and **F** were generated by file concatenation of the gated FDCs in 4 biological replicates. Graphs show mean and error bars show SD. For **B, D and E** P-value by two-tailed unpaired T test with Welch’s correction.

### *Ngfr* KO mice have spontaneous germinal center formation

To further explore the role of NGFR in this context, we characterized the SLOs of a *Ngfr* KO mouse model.^28^ First, we looked for anatomic abnormalities in the LNs of *Ngfr* KO mice. We found that *Ngfr* KO LNs were significantly larger compared to LNs from WT animals. Differences were particularly evident in popliteal (PO) LNs (**Figure 2A**), whose weight was increased about 5-fold on average (**Figure 2B**) and similar data were found for other LNs (**Supplementary figure 1A, B**). Due to the observed LN hyperplasia, we next compared immunohistochemical (IHC) staining for CD45R/B220, Ki67, and BCL6 in LNs from WT and *Ngfr* KO mice. *Ngfr* KO mice exhibited a substantial hyperplasia of the B-cell compartment based on the expression of CD45R/B220^+^ (**Figure 2C**). Moreover, BCL6^+^ GCs found in *Ngfr* KO LNs (**Figure 2C and supplementary figure 1C**) also stained with the Ki67 proliferation marker, unlike WT mice (**Figure 2C**).

**Figure 2:**
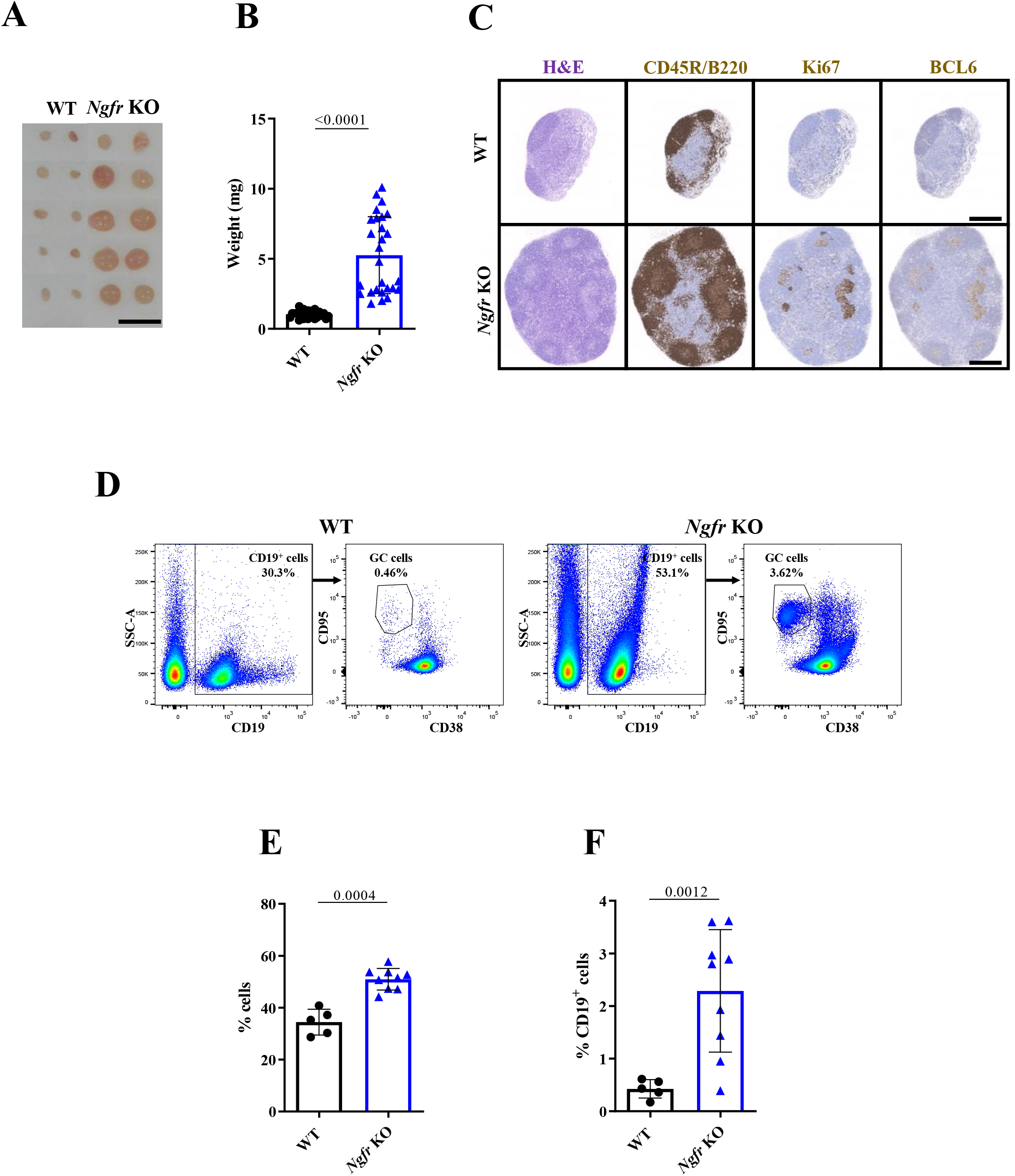
NGFR absence in the LN microenvironment leads to GC hyperplasia. **A**) Representative image of PO LN size in WT and *Ngfr* KO mice. Scale bar, 5mm. **B**) PO LN weight in WT and *Ngfr* KO mice. n=28 from 3 independent experiments. **C**) Representative H&E and immunohistochemistry of CD45R/B220, Ki67 and BCL6 in PO LNs of WT (upper panels) and *Ngfr* KO (lower panels). Scale bar, 400µm. **D**) Gating strategy for CD19^+^ and GC cells and representative samples of PO LN from WT (left panels) and *Ngfr* KO (right panels) mice. **E**) Quantification by flow cytometry of PO LN CD19^+^ cells in WT and *Ngfr* KO mice. n=5 from 1 experiment for WT and n=9 from 2 independent experiments in *Ngfr* KO mice. **F**) Quantification by flow cytometry of PO LN GC cells in WT and *Ngfr* KO mice. n=5 from 1 experiment for WT and n=9 from 2 independent experiments in *Ngfr* KO mice. Graphs show mean and error bars show SD. For **B**, P-value by two-tailed Mann-Whitney test. For **E** and **F** P-value by two-tailed unpaired T test with Welch’s correction.

We next performed flow cytometry analysis of GC B-cells in the LNs (**Figure 2D**). In agreement with data obtained by IHC, we observed that the total CD19^+^ B-cell compartment was increased in *Ngfr* KO compared to WT LNs (**Figure 2D, E and supplementary figure 1D**). Furthermore, the proportion of CD19^+^CD95^+^CD38^lo^ GC B cells was significantly increased (**Figure 2D, F and Supplementary figure 1E**). Altogether, these data demonstrate that the loss of NGFR is sufficient to trigger spontaneous GC B-cell activation within LNs.

### FDCs are the main population altered in *Ngfr* KO mice

We next explored whether the spontaneous B-cell activation and GC hyperplasia in *Ngfr* KO mice were caused by B-cell intrinsic or extrinsic mechanisms.

To explore a possible implication of NGFR in early B-cell development, we analyzed NGFR expression in different steps of bone marrow (BM) B-cell precursors maturation (Pro B cells, Pre BI cells, Pre BII cells and immature B cells)^29^ (**Supplementary Figure 2A**). Again, we found that, on average, the frequency of NGFR^hi^ cells was below 3% in all the subsets, and only immature B cells exhibited a slight shift in NGFR mean fluorescence (**Supplementary figure 2B and C**). Of note, no imbalance was found in any of those populations when comparing *Ngfr* WT and KO and BM (**Supplementary figure 2**).

Flow cytometry analysis also revealed that NGFR expression in the majority of total B cells (CD19^+^) in WT immunized LNs was negative or low, displaying a shift in NGFR fluorescence compared to *Ngfr* KO B cells, similar to the one found in immature B cells in the BM (**Supplementary figure 2E, F upper panels**). Nonetheless, we identified a minor subpopulation of approximately 20% of the CD19^+^CD95^+^CD38^lo^ GC B cells expressing dim levels of NGFR (**Supplementary figure 2E, F lower panels**).

Next, to understand if the lack of NGFR expression in GC B cells was responsible for the LN hyperplasic phenotype found in *Ngfr* KO mice, we performed BM transplantation experiments using donor BM cells from WT and *Ngfr* KO mice and sublethally irradiated WT and *Ngfr* KO recipient mice (**Figure 3A)**. We found that LN enlargement was independent on the BM origin and only occurred when the transplanted host was *Ngfr* KO (**Figure 3B**). Accordingly, flow cytometry analysis demonstrated that the B-cell compartment was significantly expanded only in *Ngfr* KO hosts independently on the BM origin (**Figure 3 C**). Importantly, while WT hosts had barely detectable GC B cells (less than 0.2% of all LN B cells), *Ngfr* KO host mice showed about 4 to 5% of GC B cells within the CD19^+^ B cell population (**Figure 3D**). These results demonstrate that GC hyperplasia, which only occurred in *Ngfr* KO mice LNs, relies on the host microenvironment rather than on the transplanted immune cells, strongly suggesting the involvement of non-hematopoietic cell populations.

**Fig 3:**
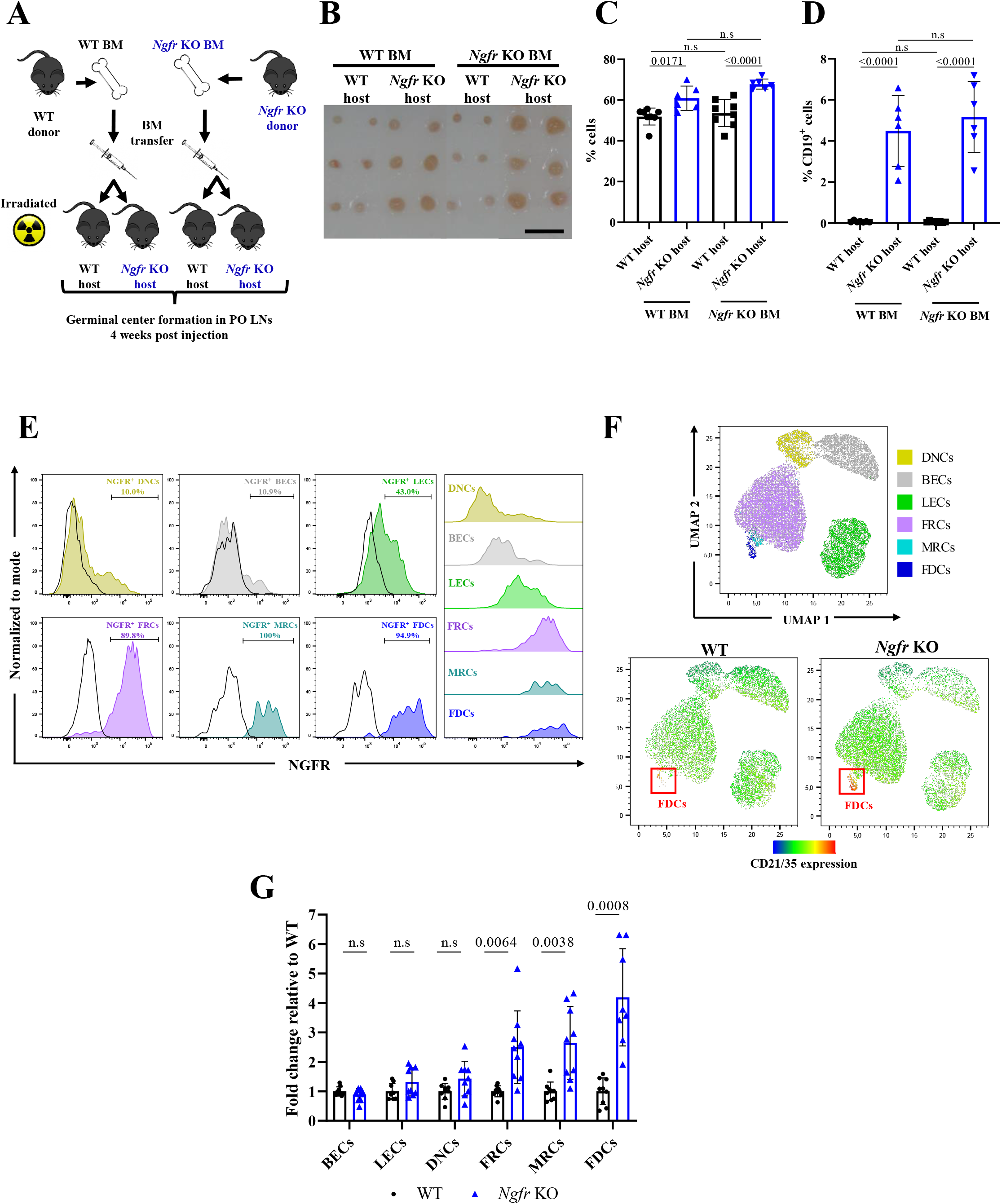
GC hyperplasia is driven by *Ngfr* KO stromal cells and FDCs are the main altered population. **A**) Experimental set-up for BM transference in irradiated WT and *Ngfr* KO mice. **B**) Representative image of PO LN size in irradiated WT and *Ngfr* KO mice 4 weeks upon BM transplant as indicated in **A.** Scale bar, 5mm. **C**) Quantification by flow cytometry of PO LN CD19^+^ cells in WT and *Ngfr* KO irradiated mice as indicated in **A**. n=8 from 2 independent experiments for WT host and n=6 from 2 independent experiments for *Ngfr* KO host mice. **D**) Quantification by flow cytometry of PO LN GC cells in WT and *Ngfr* KO irradiated mice as indicated in **A**. n=8 from 2 independent experiments for WT host (an outlier was removed from the WT BM-WT host group) and n=6 from 2 independent experiments for *Ngfr* KO host mice. **E**) Histograms comparing NGFR fluorescence in stromal populations of WT LNs. Black lines represent NGFR fluorescence in *Ngfr* KO stromal cells used as negative CTL **F**) UMAP of the stromal populations gated in **Supplementary figure 3C** (upper panel). Equal number of CD45^−^ events from 5 WT and 5 *Ngfr* KO samples were merged. Lower panels show the distribution of the intensity of CD21/35 in the UMAP separating WT and *Ngfr* KO events. **G**) Flow cytometry analysis of the different subsets of stromal populations gated in **Supplementary figure 3C**. Frequency of total cells is relative to the WT values and expressed in fold change for each population. n=9 from 2 independent experiments (an outlier was removed from the *Ngfr* KO FDCs group). Graphs show mean and error bars show SD. For **C and D** P-value by one-way ANOVA analysis and Tukey’s multiple comparisons test. For **G** P-value by two-tailed unpaired T test with Welch’s correction (except in LECs where Mann-Whitney test was performed).

To identify the non-hematopoietic cell subsets affected by *Ngfr* KO, we characterized NGFR expression in LN cells from WT mice. IHC analysis showed that most of the NGFR^+^ cells exhibited an elongated morphology and an interconnected distribution mainly confined to the cortical and B-cell areas (**Supplementary Figure 3A**). Flow cytometry analysis showed that NGFR was expressed approximately in 50% of non-hematopoietic CD45^−^ cells and only in 0.5% of the CD45^+^ hematopoietic cells (**Supplementary Figure 3B**). Moreover, analysis of CD45^−^ cell subsets (**Supplementary Figure 3C)** highlighted a dim expression of NGFR on CD31^+^PDPN^+^ lymphatic endothelial cells (LEC) and a high expression on the 3 main PDPN^+^ LSC subsets including MAdCAM-1^−^ CD21/35^−^ FRCs, MAdCAM-1^+^ CD21/35^−^ MRCs, and CD21/35^+^ FDCs **(Figure 3E and Supplementary Figure D).**

We next studied the effect of the *Ngfr* KO in LN LSC populations from naïve mice. Total non-hematopoietic CD45^−^ cells were enriched in *Ngfr* KO compared to WT mice (**Supplementary figure 3E**). We applied a dimensionality reduction approach based on the uniform manifold approximation and projection (UMAP)^30^ of our flow cytometry data obtained from both WT and *Ngfr* KO stromal cells (**Figure 3F**). We found that the main population altered in *Ngfr* KO mice corresponded to FDCs based on the expression of the canonical FDC marker CD21/35. We observed that FDCs were scarce in WT LNs while largely increased in *Ngfr* KO mice (**Figure 3F**). More precisely, whereas BECs, LECs and CD31^−^PDPN^−^ doble negative cells (DNCs) were not affected in *Ngfr* KO mice, FRCs, MRCs, and FDCs were significantly expanded (**Figure 3G).** Overall, these data support that, rather than having B-cell intrinsic effects, *Ngfr* KO affects mainly the LN LSC compartment, leading to an expansion of LSCs, in particular FDCs.

### NGFR deficiency promotes FDCs activation and up-regulates B cell activating signatures

Given that FDCs were the most expanded stromal population in *Ngfr* KO LNs and taking into account their well stablished roles in the GC physiology^14,15,18^ we wondered if FDC activation could be also affected in KO mice, contributing to B cell expansion.

We sorted CD45^−^CD31^−^PDPN^+^CD21/CD35^+^ LN FDCs from naïve WT and *Ngfr* KO mice and performed RNA sequencing (RNAseq) using SMART-Seq RNAseq technology. Principal component analysis (PCA) showed that samples clustered based on their NGFR status (**Figure 4A**). We next identified the differentially expressed genes (DEGs) between the WT and *Ngfr* KO samples. We found that 290 genes were significantly up-regulated and 157 genes down-regulated in *Ngfr* KO FDCs compared to their WT counterpart (**Figure 4B and Supplementary Table 1**).

**Figure 4:**
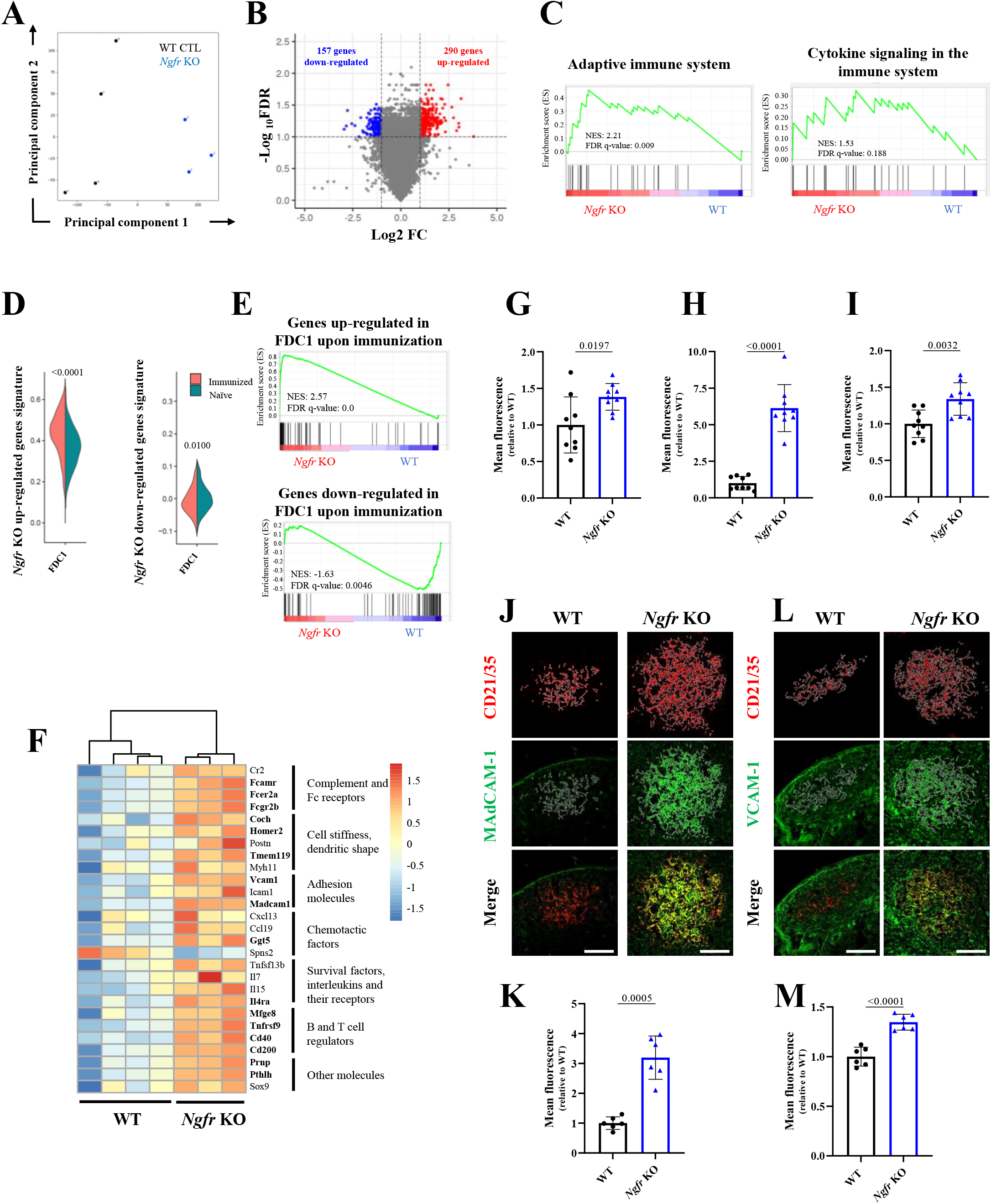
NGFR deficiency promotes FDCs activation and up-regulates B cell activating signatures. **A**) Principal component analysis of the samples included in the RNAseq analysis. n=4 for WT and n=3 for *Ngfr* KO mice. **B**) Volcano plot showing gene distribution upon differential expression analysis from *Ngfr* KO mice vs WT. Colored dots represent genes with FDR<0.1 and FC>1 (red) or <-1 (blue), see Materials and Methods section for more details. **C**) Enrichment plot from the Reactome adaptive immune system signature (right panel), and cytokine signaling in the immune system (left panel) upon GSEA with the DEGs obtained from the RNAseq analysis in FDCs. **D**) Violin plots showing the percentage of identity for the DEG signatures in *Ngfr* KO mice vs WT FDCs compared with FDCs1 (LZ-FDCs) subset found by Pikor NB et al.^26^ in immunized (orange) and naïve (green) LNs. **E**) Enrichment plots of GSEA of the differentially expressed genes between FDCs1 immunized and naïve LNs in Pikor NB et al compared to the DEG signatures in *Ngfr* KO mice vs WT FDCs. **F**) Heatmap showing the differential expression of the RNAseq analysis of *Ngfr* KO mice vs WT FDCs in a selection of the main genes involved in FDCs biology. Color code shows relative expression standardized by line. Genes in bold are among the DEGs found in the comparison between *Ngfr* KO mice vs WT FDCs (**Supplementary table 1)**. **G, H, I**) Quantification of flow cytometry data obtained from the analysis of **G**) CD21/35; **H**) MAdCAM-1 and **I**) VCAM-1 mean fluorescence in FDCs from WT and *Ngfr* KO PO LNs. n=9 from 2 independent experiments. **J, L**) Representative images from the analysis of CD21/35 (red), **J**) MAdCAM-1 (green) and **L**) VCAM-1 (green) by IF in WT and *Ngfr* KO FDCs. White line shows the area selected for quantification based on CD21/35 expression. Scale bar, 100µm. **K, M**) Quantification of **K**) MAdCAM-1 and **M**) VCAM-1 mean fluorescence in FDCs by IF in WT and *Ngfr* KO PO LNs. Up to 3 individual follicles per LN were analyzed when possible and average values were plotted. n=6 from 1 experiment. Graphs show mean and error bars show SD. For **D**, **G, H, I, K** and **M** by P-value by two tailed unpaired T test with Welch’s.

Gene Set Enrichment Analysis (GSEA) using Reactome database revealed enrichment for adaptive immune system, and cytokine signaling in the immune system pathways in *Ngfr* KO FDCs (**Figure 4C**). In agreement, genes involved in the modulation of the GC reaction, including *Fcer2, Tnfrsf9, Madcam1, Cd40, Il4r, Vcam1* or *Cd80* were all up-regulated in the *Ngfr* KO FDCs (**Supplementary Table 1**).

We next compared our *Ngfr* KO FDC signature (**Supplementary Table 1**) with previously published single cell RNAseq datasets from specific subsets of murine LSCs non-endothelial LN stromal cells.^26,31^ An initial comparison with data from whole purified LSCs showed that up-regulated genes in *Ngfr* KO FDCs were significantly enriched in the FDC subset^31^ (**Supplementary figure 4A, B, C)**. Additionally, we generated heatmaps from the top 10 DEG for each of the 9 subsets identified by Rodda LB et al. and found that the 10 top genes defining the FDC subset (including *Cr2, Fcgr2b* or *Fcer2a*)^31^ were all up-regulated in *Ngfr* KO FDCs (**Supplementary figure 4D**), supporting that *Ngfr* KO FDCs up-regulated genes correlate with the FDC canonical signature. We then compared our signature with a published scRNAseq dataset restricted to mouse B-cell interacting stromal cells identified as CXCL13-expressing cells LN.^26^ Among the 7 distinct stromal cell subsets identified in this study,^26^ the up-regulated genes in *Ngfr* KO FDCs were found to be enriched in the FDC1 cluster, corresponding to LZ-FDCs, compared to DZ-FDCs, MRCs, and FRC subsets (**Supplementary figure 3E, F, G**). Additionally, since this dataset contains gene expression profiles of LSC from naïve and immunized animals, we studied the correlation between our *Ngfr* KO FDC gene signature and the genes induced in LZ-FDCs upon immunization. Interestingly, we found that up-regulated genes in FDCs from our *Ngfr* KO mice were enriched in LZ-FDCs from immunized versus naïve WT mice^26^ (**Figure 4D**). We next extracted the 174 DEG between naïve and immunized LZ-FDCs^26^ obtaining 84 up-regulated and 90 down-regulated genes (**Supplementary Table 2**). GSEA showed that up-regulated genes upon immunization correlated with the most up-regulated genes in *Ngfr* KO FDCs while the opposite results were found for the down-regulated signature (**Figure 4E**).

We generated a curated list of genes defining FDC markers modulated in the *Ngfr* KO FDCs that were also modulated upon inflammation in WT FDCs (**Figure 4F and Supplementary Table 3**) and validated the expression of some of these markers by flow cytometry. We found that CD21/35, MAdCAM-1, and VCAM-1 were also up-regulated at the protein level in *Ngfr* KO FDCs compared to WT FDCs (**Figure 3G, H, I**). Moreover, we analyzed the expression of MAdCAM-1 (**Figure 3J, K**) and VCAM-1 (**Figure 3L, M**) in situ in CD21/35^+^ cells by immunohistofluorescence, showing that these markers were up-regulated about 3 and 1.5 fold respectively (**Figure 3J, L**). Overall, these data show that CD21/35^+^ LSCs that spontaneously formed in naïve *Ngfr* KO mice, display an activated phenotype with similar transcriptomic and phenotypic profiles than LZ-FDCs from immunized mice.

### *Ngfr* KO mice show aberrant GC structure and functionality

We compared GC formation in immunized WT mice and naïve *Ngfr* KO mice. Analysis of BCL6 and CD21/CD35 expression by IHC showed that *Ngfr* KO mice have aberrant GC localization within LN medulla compared to WT mice (**Figure 5A)**. In addition, some clusters of FDCs in the medulla did not co-localize with BCL6 expression (**Figure 5A, lower panels and data not shown**).

**Figure 5:**
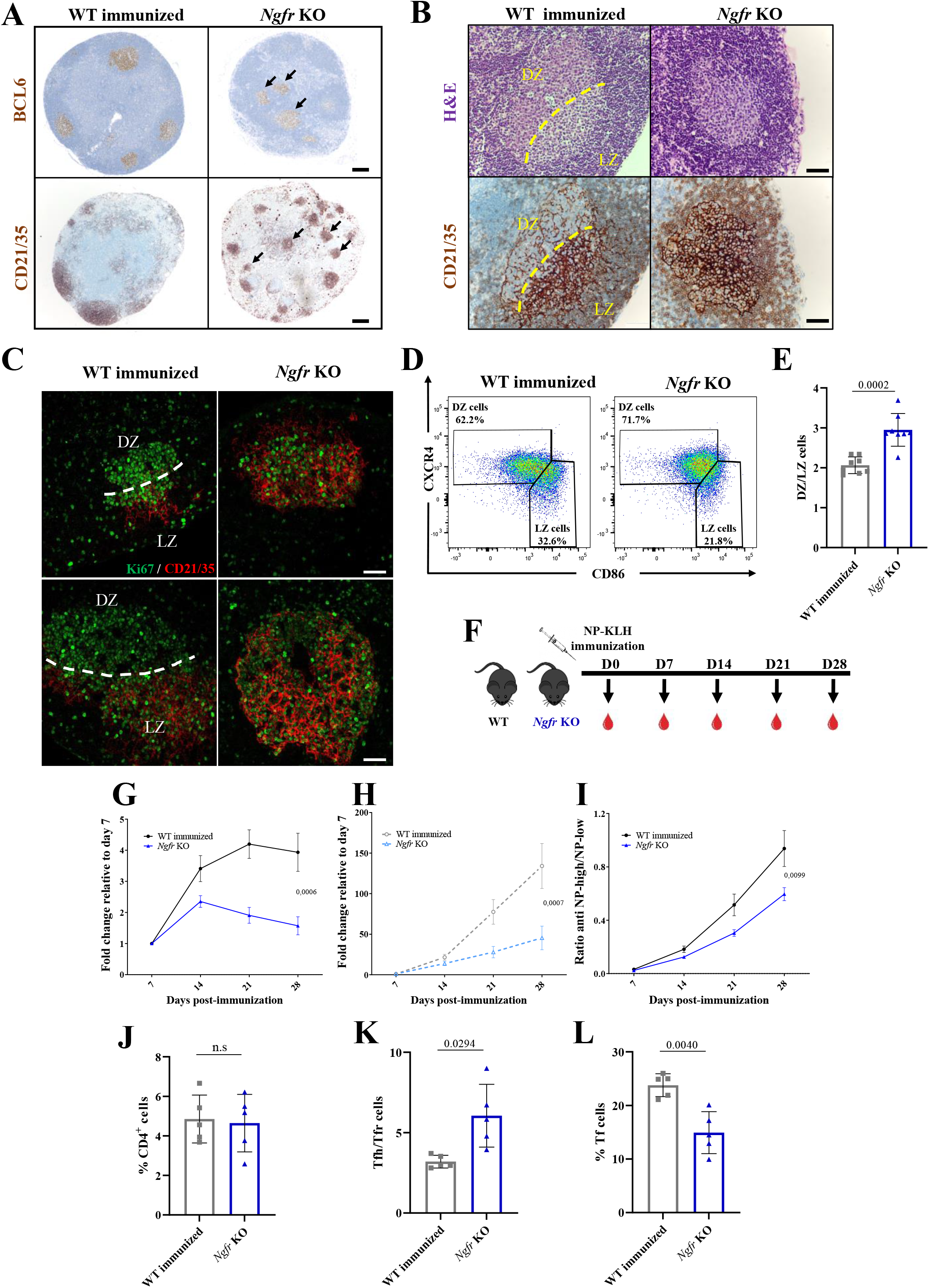
NGFR absence in FDCs alters GC functionality and B cell phenotype. **A**) Representative images of BCL6 and CD21/35 staining in PO LNs from WT immunized and *Ngfr* KO mice. Black arrows show formation of misplaced GCs. Scale bar, 200µm. **B**) Representative images of H&E (upper panels) and CD21/35 staining (lower panels) in GCs from WT immunized and *Ngfr* KO PO LNs. Yellow-dashed line divide GC DZ and LZ. Scale bar, 50µm. **C**) Representative images of Ki67 (green) and CD21/35 (red) staining in GCs from WT immunized and *Ngfr* KO PO LNs. White-dashed line divide GC DZ and LZ. Scale bar, 50µm. **D**) Gating strategy used in flow cytometry analysis of centroblasts (DZ cells) and centrocytes (LZ cells). **E**) Quantification of DZ/LZ ratio based on the frequencies of centroblast and centrocytes in WT immunized and *Ngfr* KO PO LNs. n=8 from 2 independent experiment. **F**) Scheme of intrafootpad NP-KLH immunization and blood sampling in WT and *Ngfr* KO mice. **G**) Fold increase of low-affinity antibodies normalized at day 7. **H**) Fold increase of high-affinity antibodies normalized at day 7. **I**) Ratio between high- and low-affinity antibodies in serum from WT and *Ngfr* KO mice. **J**) Quantification by flow cytometry of total Tf cells (CD3^+^ CD4^+^CXCR5^+^ PD-1^+^) cells in WT immunized and *Ngfr* KO PO LNs. **K**) Quantification by flow cytometry of the ratio between Tfh (FOXP3^−^) and Tfr cells (FOXP3^+^) cells in WT immunized and *Ngfr* KO PO LNs **L**) Quantification by flow cytometry of Tfr cells in WT immunized and *Ngfr* KO PO LNs. For **G**, **H**, **I, J, K,** and **L** n=5 from 1 experiment. For **E, J, K** and **L** Graphs show mean and error bars show SD. P-value by two-tailed unpaired T test with Welch’s correction. For **G**, **H**, and **I** Graphs show mean and error bars show SEM. P-value by two-way ANOVA.

Studies of GC structure showed that *Ngfr* KO mice displayed major alterations. H&E staining highlighted classical LZ/DZ areas within GC from immunized WT mice whereas such organization was disrupted in most GCs of *Ngfr* KO mice, which homogeneously displayed a histological appearance more similar to the DZ area (**Figure 5B, upper panels**). Since FDCs are fundamental organizers of the GC structure, we analyzed CD21/35 expression (**Figure 5B, lower panels**). We observed a typical distribution of CD21/35^+^ cells in the LZ of GCs from immunized WT mice. Conversely, CD21/35^+^ cells were homogenously distributed in the GCs from naive *Ngfr* KO mice with no apparent zonation (**Figure 5B, lower panels**). To further characterize GC features in *Ngfr* KO mice, we studied GC B-cell proliferation using Ki67 staining. Proliferation was mainly restricted to CD21/35^−^ DZ area in GCs from WT immunized mice, while it was homogeneously distributed in the GCs from *Ngfr* KO mice (**Figure 5C**). We also analyzed by flow cytometry the frequency of LZ and DZ B cells in the LNs of immunized WT and naïve *Ngfr* KO mice based on the expression of CD86 and CXCR4 markers^32^ (**Figure 5D**). The DZ/LZ B-cell ratio was significantly increased in *Ngfr* KO mice compared to immunized wT mice (**Figure 5E**). These data confirm that, despite being surrounded by LZ-like FDCs, GC B cells are skewed towards a DZ phenotype in naïve *Ngfr* KO mice.

Since GC polarization and iterative transition of GC B cells through the DZ and the LZ is a key feature of these microstructures, ensuring correct B-cell selection,^18^ we evaluated if the disruption observed in *Ngfr* KO mice had an impact on the generation of high-affinity antibodies. For that purpose, we immunized WT and *Ngfr* KO mice with NP-KLH by intrafootpad injection and analyzed the generation of high- and low-affinity IgG1 antibodies against NP every 7 days during 4 weeks post-immunization (**Figure 5F**). To analyze the differences, we normalized antibody levels referred to the initial titers measured at day 7. We found that both types of antibodies showed a significantly increased production over time in WT mice. At day 28, the fold change relative to day 7 was 2.36- and 88.66-fold higher for low- and high-affinity antibodies in WT mice compared to the *Ngfr* KO respectively (**Figure 5G, H**). We established the ratio between high- and low-affinity antibody levels to evaluate the effectiveness of the affinity maturation process.^25^ This value is close to zero at initial time points and increases upon time as high-affinity clones are selected and antibodies are secreted to the bloodstream. We observed that ratios between high- and low-affinity antibodies were significantly higher in WT than in the *Ngfr* KO mice (**Figure 5I**), suggesting that the efficiency of the high-affinity humoral response in the *Ngfr* KO mice is compromised over time.

Due to the pivotal role of T follicular (Tf) cells in the affinity maturation process, we wondered if this population was altered in the *Ngfr* KO GCs, contributing to impair the humoral response. Flow cytometry analysis discarded alterations in total T cells (CD3^+^) as well as CD4^+^ and CD4^−^ T cells in *Ngfr* KO compared to WT immunized LNs (**Supplementary figure 5A, B, C, D)**. Importantly, despite total Tf cells (CD3^+^CD4^+^CXCR5^+^PD-1^+^) also remain unaltered (**Supplementary figure 5A and Figure 5J**), we observed an alteration in the ratio between the T follicular helper (Tfh) cells (CD3^+^ CD4^+^CXCR5^+^ PD-1^+^ FOXP3^−^) and the T follicular regulatory (Tfr) cells (CD3^+^ CD4^+^CXCR5^+^ PD-1^+^ FOXP3^+^) (**Figure 5K**) due to a significant decrease of the Tfr population in *Ngfr* KO LNs (**Figure 5L**).

### *Ngfr* KO boosts pathological phenotypes and decreases survival by enhancing autoimmunity in a BCL2 overexpressing context

Since the activation of GCs without appropriate B cell selection has been directly linked to the uprising of autoreactive clones,^21^ and taking into account the decreased frequency of Tfr cells in the GCs of *Ngfr* KO mice, we evaluated the effect of *Ngfr* KO in autoimmunity. We performed immunofluorescence staining to measure the presence of IgG anti-nuclear antibodies (ANAs) in the serum from 10-week-old WT and *Ngfr* KO mice. We found that *Ngfr* KO mice exhibited detectable levels of ANA IgGs at 1/50 and 1/250 dilutions (**Supplementary figure 5E, F**), demonstrating the presence of circulating autoantibodies. By contrast, none of the tested sera from control WT mice displayed Hep2 reactivity at any dilution. We also evaluated if the presence of autoantibodies in *Ngfr* KO mice could eventually contribute to the development of autoimmune diseases in aged animals. The follow up of a cohort of 7 WT and 13 *Ngfr* KO mice up to 85 weeks, revealed no clinical signs of increased mortality or autoimmune damage in the evaluated organs (including lungs, liver, kidneys and pancreas) (data not shown). Nevertheless, the generation of self-reactive antibodies is an abnormal feature indicating a dysregulated immune response in the *Ngfr* KO mice. For this reason, we decided to further study this phenotype in a context more prone to developing autoimmune lesions.

We crossed *Ngfr* KO mice with a transgenic model overexpressing the human gene *BCL2* controlled by Vav gene regulatory (VavP) sequences (abbreviated as *Bcl2* Tg).^33^ This model has been previously used to analyze autoimmune diseases *in vivo*.^34,35^ We studied the formation of GCs in 10-week-old mice. LNs from *Bcl2* Tg/*Ngfr* KO mice were enlarged compared to *Bcl2* Tg/*Ngfr* WT mice with an increased expression of BCL6 by IHC (**Supplementary figure 6A**). Flow cytometry analysis confirmed that GC B cells were increased in *Bcl2* Tg/*Ngfr* KO LNs compared to *Bcl2* Tg/*Ngfr* WT mice (**Supplementary figure 6B, C, D**), together with an increase in FDC frequency (**Supplementary figure 6E, F).**

We further analyzed the *Bcl2* Tg/*Ngfr* KO model at later stages. Previous studies with the *Bcl2* Tg model showed that about 20% of the animals need to be sacrificed before 35-40 weeks of age due to the development of autoimmune syndromes which particularly affect the kidneys of the animals, leading to the development of autoimmune-like glomerulonephritis.^33^ We studied a cohort of *Bcl2* Tg/*Ngfr* WT and *Bcl2* Tg*/Ngfr* KO mice at 35 weeks of age. We analyzed the kidneys of these mice and performed histological analysis looking for pathological alterations.^33^ We quantified the percentage of glomeruli affected by: 1) crescent formation due to Bowman’s capsule epithelium hyperplasia (a sign of advanced kidney affectation), 2) the generation of eosinophilic deposits within the glomeruli (typically found in this model when animals are affected by autoimmunity and related to the accumulation of immunoglobulins)^34,35^ (**Figure 6A, B**). We found that the percentages of glomeruli affected by Bowman’s capsule hyperplasia (**Figure 6C**) and eosinophilic deposits (**Figure 6D**) were significantly higher in the *Bcl2* Tg/*Ngfr* KO mice compared to *Bcl2* Tg/*Ngfr* WT mice (**Figure 6C, D**), suggesting that *Ngfr* KO can accelerate the appearance of autoimmune disorders in this mouse strain.

**Figure 6:**
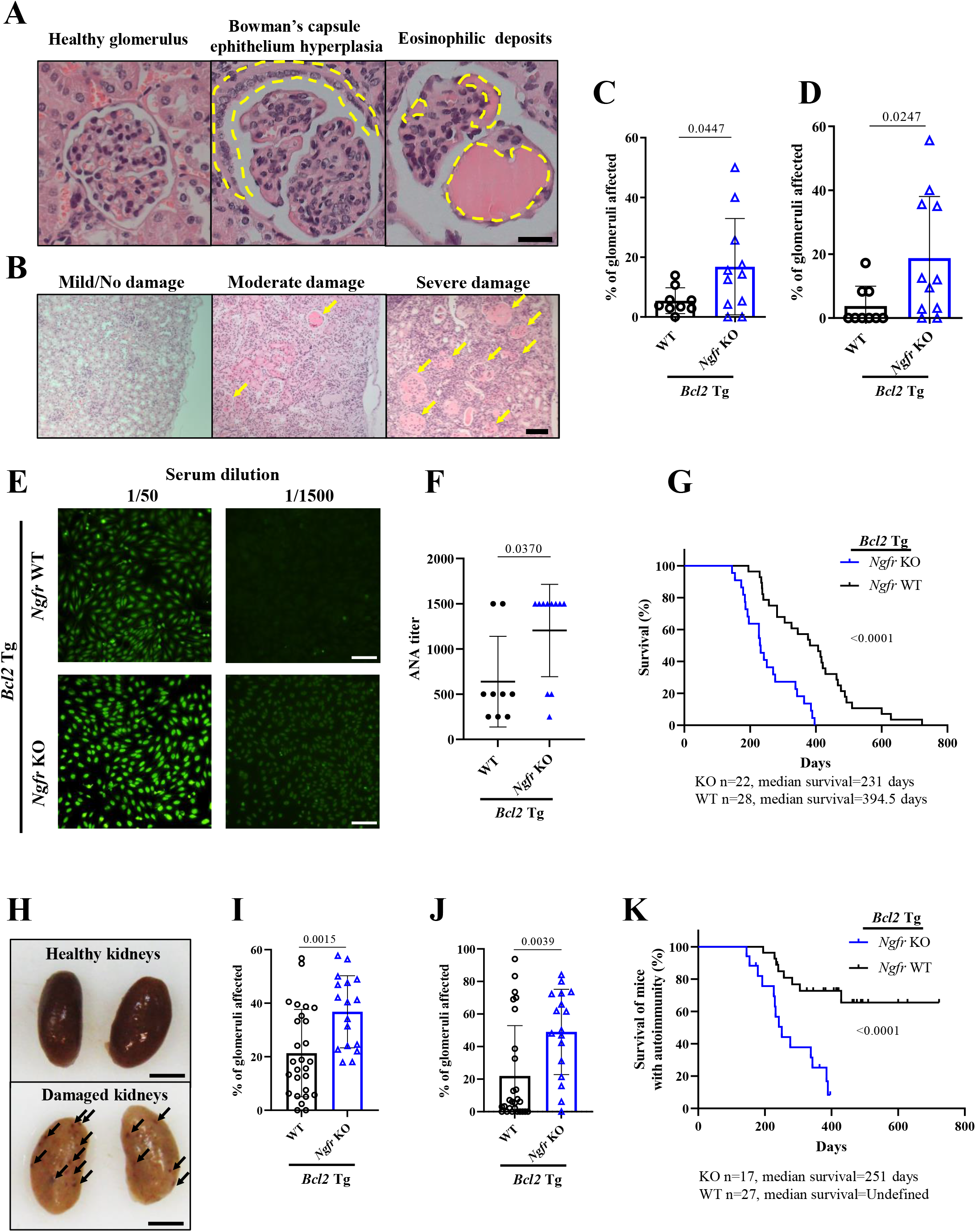
*Ngfr* KO boosts pathological phenotypes and decrease survival by enhancing autoimmunity in a BCL2 overexpressing context. **A**) Representative images from a healthy glomerulus (left panel) and the two markers studied to assess kidney damage (central and right panels). Scale bar, 2µm. **B**) Representative images of H&E from kidneys with mild, moderate or severe kidney damage. Scale bar, 10µm. **C**) Quantification of the percentage of glomeruli affected by Bowman’s capsule epithelium hyperplasia in the 35-week-old cohort. **D**) Quantification of the percentage of glomeruli affected by eosinophilic material deposits in the 35-week-old cohort. **E**) Representative images of IF analyzing ANAs using serum from 35-week-old *Bcl2* Tg/*Ngfr* WT and KO mice incubated at indicated dilutions in Hep-2 cells slides. **F**) Semi-quantitative measurement of ANAs titer in *Bcl2* Tg/*Ngfr* WT and KO 35-week-old mice serums. n=9 *Bcl2* Tg/*Ngfr* WT n*=*11 *Bcl2* Tg/*Ngfr* KO mice. **G**) Survival curve of *Bcl2* Tg/*Ngfr* WT and KO mice. **H**) Representative image from healthy (upper panel) and damaged (lower panel) kidneys from *Bcl2* Tg mice. Black arrows denote areas of macroscopic hemorrhagic foci. **I**) Quantification of the percentage of glomeruli affected by Bowman’s capsule epithelium hyperplasia in the survival cohort**. J**) Quantification of the percentage of glomeruli affected by eosinophilic material deposits in the survival cohort. For **I** and **J** n=27 *Bcl2* Tg/*Ngfr* WT and n=17 *Bcl2* Tg/*Ngfr* KO. **K**) Survival curve of *Bcl2* Tg/*Ngfr* WT and KO mice affected with autoimmunity (presenting more than 20% of the glomeruli with signs of eosinophilic deposits). Graphs show mean and error bars show SD. For **C and I** P-value by two-tailed unpaired T test with Welch’s correction. For **D, F and J** P-value by two-tailed Mann-Whitney test. For **G** and **K** P-value by Log-rank (Mantel-Cox) test.

We also assessed the levels of ANAs in these cohorts of 35-week-old animals. ANA titers in the sera of *Bcl2* Tg/*Ngfr* KO mice were significantly higher than *Bcl2* Tg/*Ngfr* WT mice with most samples exhibiting homogeneous nuclear staining even at the highest dilution tested (1/1500) (**Figure 6E, F**).

To further evaluate if *Ngfr* KO could lead to an increased incidence of severe autoimmune phenotypes, we decided to study the long-term effects of the *Ngfr* KO in this pathology. We established additional cohorts of *Bcl2* Tg/*Ngfr* WT and *Bcl2* Tg/*Ngfr* KO mice to study their long-term evolution and overall survival (OS). We found significant changes in OS with a median survival of 231 days for the *Bcl2* Tg/*Ngfr* KO mice compared to 394.5 days in the *Bcl2* Tg/*Ngfr* WT group (**Figure 6G**). We looked for signs of damage in the kidneys of these cohorts of aged animals. Macroscopic hemorrhagic foci and loss of color could be detected in most of the early dying mice (**Figure 6H**), suggesting severe kidney damage. We performed additional histological analysis to assess Bowman’s epithelium hyperproliferation and the presence of eosinophilic deposits. The majority of the *Bcl2* Tg/*Ngfr* KO mice in this end-point cohort showed severe kidney damage with more than 30% of their glomeruli affected by any of the quantified lesions or both concomitantly, whereas *Bcl2* Tg*/Ngfr* KO mice exhibited less severe kidney lesions (**Figure 6I, J**). Of note, the phenotype found in the kidneys of the *Bcl2* Tg/*Ngfr* WT mice was predominantly mild or moderate in alive mice after 35 weeks compared to those that died before 35 weeks (approximately 20% of the cohort).

Importantly, IHC analysis of eosinophilic deposits found in the kidney of *Bcl2* Tg/*Ngfr* KO mice was positive for IgG (data not shown), suggesting that autoantibodies generation in these mice might be the cause of the observed tissue damage. For this reason, we then established the presence of eosinophilic deposits in more than 20% of the glomeruli as a criterion to stratify mice affected by autoimmunity at endpoint. Having in mind this criterion, we found that 30% of the *Bcl2* Tg/*Ngfr* WT mice died with signs of autoimmunity compared to more than 80% of the *Bcl2* Tg/*Ngfr* KO animals (**Figure 6K**).

## Discussion

Despite NGFR expression in immune cells and lymphoid organs has been reported,^36^ its role in the regulation of LN functionality is still unclear. In this work, we analyzed the role of NGFR in LNs using a NGFR KO model. We found that *Ngfr* KO mice showed LN hyperplasia associated with expansion of the GC B-cell compartment. Our analysis by flow cytometry showed that NGFR is barely expressed on the surface of BM and LN B cells, and no significant differences in the distribution of B cell precursor populations were observed in *Ngfr* KO BMs. Our findings align with previous studies indicating that NGFR expression in murine B cells remains low but it is up-regulated in antibody-secreting plasma cells of lupus-prone mice.^37^ Of note, in this study, *Ngfr* KO in B cells attenuates autoimmune disease progression and negatively impacted the generation of B cells in the spleen. Importantly, our BM chimera experiments revealed that the hyperplastic GC phenotype observed was not primarily attributed to *Ngfr* deletion in lymphoid cells. Interestingly, LN hyperplasia only occurred in *Ngfr* KO hosts, regardless of *Ngfr* expression in hematopoietic cells. These findings indicate that the activation of GCs is dependent on *Ngfr* KO in radioresistant stromal cells of the LN microenvironment.

Our analysis showed that NGFR was widely expressed in LECs and in different subsets of LSCs including FRCs, MRCs, and FDCs. Although the expression of NGFR in some of these populations has been already reported,^12,38,39^ its function remains largely unexplored. We identified FDCs as the most expanded LSC subset in the *Ngfr* KO LNs, consistent with the GC hyperplasia observed in these mice.

FDCs are key players modulating GC physiology.^14–16^ Despite our knowledge about FDC biology has slowly increased during the last decades, there are still many aspects about their behavior that remain elusive due to their low frequency (less than 1% of all LN stroma cells)^40^, limitations in their purification, and impossibility to maintain them in culture as functional cells.^41–44^ Moreover, study of human FDCs has been mainly performed in pathological conditions, including lymphoid neoplasia or chronically inflamed resected tonsils^45,46^. Our analysis comparing WT FDCs in naïve and immunized mice revealed that NGFR is down-regulated upon FDC activation. Furthermore, we demonstrated that NGFR is essential in restricting FDC activation in the absence of inflammation. *Ngfr* KO FDCs spontaneously developed an activated phenotype, similar to the one described during immune responses,^26^ with the modulation of genes regulating FDC maturation as LZ-FDCs.^14,26,47–54^ Several questions remain open in this regard, such as the potential utility of NGFR as a marker for stratifying distinct subsets of FDCs or assessing their activation status, as well as its biological significance for each subset.

Tumor necrosis factor (TNF)-α and lymphotoxin (LT)-α1β2 are the two non-redundant factors required for FDC maturation from stromal precursors, both within secondary lymphoid organs and in ectopic tertiary lymphoid structures.^14,55,56^ Few other stimuli, like retinoic acid or toll-like receptor ligands, have been also reported to contribute to FDC differentiation and function.^57,58^ Of note, one of the main molecular cues regulated by TNF and LT in FDCs is the NF-κB pathway.^59,60^ They can induce I-kappa-B kinase 2 (IKK2) phosphorylation, triggering this way the classical NF-κB signaling cascade. This promotes the up-regulation of adhesion molecules as ICAM-1 and VCAM-1 in FDCs, while IKK2 mutant mice fails triggering their expression, leading to the formation of dysfunctional GCs.^14^ It is worth noting that NF-κB is one of the most relevant pathways modulated by NGFR.^61^ In this specific context, a crucial question that remains unanswered is the precise mechanism by which NGFR signaling modulates FDC differentiation and maturation into LZ-FDC.

We found major structural alterations in the *Ngfr* KO GCs with a marked loss of zonation and an aberrant distribution of the FDCs network. GC zonation is a feature conserved across species,^18^ suggesting an important role in evolution. However, the precise molecular mechanisms that drive the maturation and polarization of naïve FDCs into GC FDCs are still not fully comprehended. Nevertheless, it is believed that the interaction between FDCs and GC B cells, facilitated by chemotactic factors such as CXCL12 and CXCL13, plays a pivotal role in this process.^32,62^ In fact, the generation of chemotactic gradients in SLOs is crucial for the organization of these organs and ensures the proper distribution of the different cell populations.^63^ Recent studies have demonstrated that mouse spleen contains a specific number of FDC clusters which serves as GC niches. These niches could be filled by GC B cells in response to inflammation or immunization without increasing the number of FDC clusters.^64^ Our study has unveiled new avenues for investigating the influence of NGFR and the potential role of neurotrophins in FDC polarization and positioning.

It is important to acknowledge that our study utilized a whole-body constitutive KO model,^28^ which presents certain limitations when interpreting the results. We could not exclude if the observed phenotype is exclusively caused due to the deficiency of NGFR in LZ-FDCs or if there are contributions from other cell populations in KO mice. In this context, one possible explanation is that the maturation and expansion of the DZ-FDCs are impaired in the *Ngfr* KO LNs, resulting in the available space within the GC being occupied by LZ-FDCs. Further investigations employing conditional models to selectively deplete *Ngfr* from specific stromal populations, including different subpopulations of FDCs, will be crucial in elucidating this matter.

Additionally, we found detectable levels of self-reactive ANAs in the serum of *Ngfr* KO mice. There is evidence in the literature supporting that alterations in GCs could contribute to the expansion of self-reactive clones.^21,65,66^ In this context, FDCs play a dual role since they participate in the selection of high-affinity B cell clones, optimizing exposition of external antigens and survival factors,^18,67^ but they can also expose self-antigens,^21,68^ which may contribute to the development of autoimmunity. In our model, we observed a decreased frequency of Tfr cells within GCs of *Ngfr* KO LNs compared to immunized WT mice. This finding suggests that the reduction in Tfr cells could potentially contribute to the survival of self-reactive clones and the development of autoimmune syndromes as previously reported.^69,70^ FDCs are involved in the recruitment of Tf cells to the GCs *via* CXCL13 secretion,^14^ and their positioning within the GC has been demonstrated to be altered when polarization is lost, affecting humoral immunity.^26^ Taking into account the observed phenotype in *Ngfr* KO GCs, it is plausible to speculate that disrupted structure may also result in abnormal chemokine gradients, potentially affecting the populations of Tf lymphocytes. Nevertheless, further data are required to draw conclusion regarding whether the recruitment or functionality of Tfr cells are affected due to the absence of NGFR in FDCs.

Despite observing detectable levels of autoantibodies in the serum of 10-week-old *Ngfr* KO mice, we did not find any impact in autoimmunity in this mouse model. This may be explained attending to the multistep nature of autoimmune diseases, which normally require to bypass sequential tolerance checkpoints to develop harmful manifestations.^71^ Interestingly, a combination of *Ngfr* KO model with *Bcl2* overexpressing mice,^33^ led to decreased OS due to a higher incidence of severe lupus-like glomerulonephritis. *Bcl2* overexpressing mice have been extensively used to analyze autoimmune diseases *in vivo*.^34,35^ In this model we observed that *Ngfr* KO led to a significant increase in GC formation, FDCs frequency and subsequent aggravation of autoantibody production. Of note, the VaV-*Bcl2* model was originally developed to replicate the progression of follicular lymphoma,^33^ a chronic and incurable lymphoproliferative disease.^72,73^ Interestingly, our findings revealed that the shortened lifespan of *Bcl2* Tg/*Ngfr* KO mice, attributed to an increased incidence of severe autoimmune syndromes, hindered the development of high-grade follicular lymphomas. Consequently, the applicability of this model for studying the specific pathology of high-grade follicular lymphomas was limited.

There are several studies supporting the relevance of neurotrophin signaling in autoimmune diseases, including rheumatoid arthritis, multiple sclerosis and SLE.^74,75^ Regarding NGFR, while conditional deletion of this gene in B cells had a beneficial effect attenuating the symptoms of SLE in mice,^37^ induction of experimental autoimmune encephalomyelitis in the same model that the one used here, increased the severity and lethality of the disease.^76,77^ These studies together with ours illustrate the complex and pleiotropic nature of NGFR and the neurotrophin signaling network, suggesting that the role for NGFR in the regulation of autoimmune disorders will be different in LSCs compared to B cells. The involvement of BDNF has been extensively investigated in B cells, primarily characterized as a promoter of B cell survival through the activation of the tropomyosin receptor kinase B (TrkB) signaling.^78–80^ Moreover, a recent study demonstrated that pro-BDNF can also exacerbate autoimmunity in SLE interacting with NGFR.^37^ On the contrary, how neurotrophins specifically modulate FDC physiology remains largely unexplored. Only NGFR and the high-affinity NGF receptor TrkA have been identified in FDCs, suggesting that NGF will likely play a relevant role in this stromal population.^31,81,82^ In fact, adding supplementary NGF to FACS isolated FDCs favored its culture and the maintenance of a dendritic phenotype.^83^ Exploring the interaction between these receptors and their respective ligands, not only in B cells but particularly in FDCs throughout various stages of the GC reaction, holds significant research interest. Some reports provide evidence for the reciprocal inhibition of NGFR and TrkA expression depending on the context.^84^ This concept is consistent with recent models proposing that the transmembrane and/or intracellular domains of NGFR interact with TrkA, facilitating conformational changes with allosteric effects that modulate the affinity and specificity of TrkA for NGF.^85^ Further research is necessary to understand if the NGFR:TrkA axis is directly involved in FDC maturation or activation. In this context, our findings show that the activation of FDCs can induce alterations in NGFR expression, suggesting that FDCs might be responsive to changes in neurotrophin levels associated with both physiological and pathological scenarios.

Collectively, our data provide compelling evidence supporting the involvement of NGFR in the regulation of FDCs activation, functionality, and preserving immune tolerance. The loss of NGFR expression leads to the appearance of self-reactive clones, promoting the generation of autoantibodies which, together with coexisting alterations (*e.g.,* BCL2 overexpression), can contribute to the development of autoimmune pathology. Our study unveils a potential origin of B-cell mediated autoimmunity from stromal anomalies and a dysregulated B-cell/ FDC cross-talk. These findings prompt further exploration of whether similar anomalies in cross-talk may contribute to autoimmunity in patients. Additionally, our data suggest that modulation of NGFR expression could be a potential strategy for modulating the severity of autoimmune syndromes. Moreover, analysis of NGFR expression in FDCs in autoimmune disorders holds promise as a potential biomarker for monitoring the progression of these pathologies.

## Materials and methods

### 1) Cellular biology approaches

#### 1.1) Flow Cytometry

LN single cell suspensions were obtained as previously described by Fletcher, A. L. et al.^86^ LNs were incubated in 2 mL of a freshly prepared digestion enzyme mix containing 0.8 mg/ml Dispase II (Roche), 0.2 mg/ml Collagenase P (Roche) and 0.1 mg/ml DNase I (Roche) dissolved in RPMI-1640 (Sigma). Tubes were incubated at 37°C in a water bath gently inverting every 5 min. After 20 min, LNs were disrupted by pipetting up and down. The largest fragments were allowed to settle and the soluble fraction was removed and transferred to 10 mL of ice-cold FACS buffer (PBS, 5 mM EDTA, 0,1% BSA). Another mL of the digestion mix was added to the fragments for another 10 minutes of digestion in the water bath. The content was centrifuged 4 minutes at 300 g and 4°C. Cells were resuspended and filtered through 70 µm strainers (Corning). BM cells were collected by flushing 4 mL of PBS in femur and tibiae from donor mice with a 26G needle and further disaggregated with a 19G needle, centrifuged and filtered using 70 µm strainers (Corning). For BM cells were centrifuged 4 minutes at 300 g and 4°C after being filtered. Cell pellets were treated adding ACK lysing buffer (Lonza) for 1 minute. Lysis was stopped adding 10 mL of ice-cold FACS buffer and cells were centrifuged again. All single cell suspensions were counted in an automatic cell counter Countess 3 FL (Invitrogen) and prepared for subsequent staining and analysis as detailed below.

To perform flow cytometry, the same number of cells was stained for each sample, normally 5 million cells. Single cell suspensions were incubated for 15 minutes at 4°C with anti CD16/CD32 Mouse BD Fc Block diluted 1/50 (BD) in 100 µL of FACS buffer in 96 wells round-bottom plates (Thermo Fisher Scientific). Cells were washed with 200 µL of FACS buffer and centrifuged 4 minutes at 300 g and 4°C. The pellets were resuspended in the antibodies mix prepared for each experiment and incubated for 45 minutes at 4°C in the dark. Cells were washed, centrifuged again and eventually filtered using 30 µm strainers CellTrics (Sysmex) into 5 mL polystyrene round-bottom cytometer tubes (Corning).

For intracellular staining with anti-FOXP3, the eBioscience Foxp3/transcription factor Staining Buffer Set (Invitrogen) was used following manufacturer instructions. LIVE/DEAD Fixable aqua (Thermo Fisher Scientific) was used as a viability dye.

Data were acquired on BD FACSCanto II or LSR Fortessa X-20 cytometers (BD) using FACSDiva v.9.0 software (BD). Compensation was performed using UltraComp eBeads (Invitrogen). Data were analyzed using FlowJo software v.10.8.1 (Tree Star).

The list of antibodies used for flow cytometry are detailed in **Supplementary table 4**:

#### 1.2) Immunofluorescence

For immunofluorescence analysis, tissues were harvested and fixed overnight (ON) in PBS 4% PFA (Electron Microscopy Sciences). Fixed tissues were subsequently incubated 24 hours in a solution of sucrose 15% and 30%. Tissues were afterwards embedded in Tissue-Tek O.C.T (Sakura) blocks and stored at −80°C.

10 µm sections were cut using a Leica CM1950 cryostat (Leica) and collected in Superfrost Plus microscope slides (Thermo Fisher Scientific). For immunofluorescence, the slides were placed in a moist chamber and incubated subsequently with PBS 4% PFA (Electron Microscopy Sciences); PBS 100 mM glycine (ITW), and PBS 0.3% Triton X100 (Sigma) at RT for 10 minutes each and washed 3 times with PBS for 5 minutes between every step. Tissues were blocked for 2 hours using IF solution (PBS 0.2% Triton X100, 0.05% Tween20 (Sigma)) with 10% goat serum or donkey serum (Jackson ImmunoResearch) serum and 1% AffiniPure F(ab′)2 fragment donkey anti-mouse IgG (Jackson ImmunoResearch). Primary antibodies were diluted in IF buffer at the dilutions detailed below (**Supplementary table 5**) and incubated ON at 4°C.

Primary antibodies were washed with PBS 0.05% Tween20 (Sigma) 3 times for 5 minutes, and secondary antibodies were diluted in IF buffer and incubated for 1 hour at RT and washed 3 more times with PBS 0.05% Tween20 (Sigma). Sections were incubated with PBS DAPI (5 µg/mL) (Merck) for 20 minutes and mounted using ProLong Diamond antifade mounting medium (Invitrogen).

Images were obtained using a using a TCS SP5 or a TCS SP8 X Leica confocal microscope and LAS X Software (Leica). Data were analyzed using Fiji software.^87^

The list of antibodies used for immunofluorescence are detailed in **Supplementary table 5**:

### 2) Animal assays

All experiments with mice were performed in accordance with protocols approved by the Institutional Ethics Committee for Research and Animal Welfare (CEIyBA) of the CNIO (IACUC 012-2017), the Instituto de Salud Carlos III (ISCIII, CBA 17_2017v3) and the Comunidad Autónoma de Madrid (CAM, PROEX 225/17). The *Ngfr* KO mice were purchased from The Jackson Laboratory. The *Bcl2 Tg* mice were kindly provided by Dr. Suzanne Cory (Walter and Eliza Hall Institute of Medical Research, Melbourne, Australia). Both models were backcrossed up to 8 generations with WT mice from a C57BL/6JOlaHsd background, purchased from ENVIGO. Mice were bred in house at specific pathogen-free conditions maintained under a regular 12-hour light-dark cycle in a temperature-controlled room (22 ± 1°C). Unless indicated otherwise, animals between 8-12 weeks of age were used for the experiments. Details for specific procedures are described below.

#### 2.1) Bone marrow transplant

To generate BM chimeras, 5-6-week-old mice were irradiated with wo doses of 4,5 Gy separated by a 4-hour-interval. 8 hours after the first dose, 5 million BM cells obtained from donor mice resuspended in 100µL of PBS were infused in the irradiated animals by retro-orbital injection. BM cells were collected by flushing 4mL of PBS in femur and tibiae from donor mice using a 26G needle and further disaggregated using a 19G needle. Cells were centrifuged, filtered using 70µm strainers (Corning) and counted in a Neubauer chamber. Irradiated animals were sacrificed 4 weeks after BM reconstitution and different organs were sampled for further analysis.

#### 2.2) Immunization protocols

To study GC formation in the LNs, animals were immunized by intrafootpad injection of 10μg of 4-hydroxy-3-nitrophenyl-acetyl (NP) hapten conjugated to the Keyhole limpet hemocyanin (NP-KLH) (Biosearch Technology) mixed with 10 µL of Imject™ Alum (Thermo Fisher Scientific) as an adjuvant and PBS in a total volume of 30 µL. PO LNs were sampled at 10 days for further analysis.

#### 2.3) Antibody production assays

To study the antibody production process, animals were immunized injecting NP-KLH as described above in both footpads. Blood samples were taken from the submandibular venous sinus prior to immunization and once a week after immunization during four weeks to characterize antibody levels in serum. Blood samples were centrifuged 10 minutes at 10.000 g and serum was collected and storaged at −80°C.

ELISA to detect specific anti-NP IgG1 antibodies was performed in 96-well flat bottom Nunc-Immuno^TM^ MaxiSorp^TM^ plates (Thermo Fisher Scientific). Plates were incubated ON using 10 μg/mL of NP(2,5)-BSA or NP(25)-BSA as the coating reagent kindly provided by Dr. Alejo Efeyan. Then, wells were rinsed three times with PBS 0.04% Tween-20 and blocked with PBS 0.04% Tween-20, 2% BSA for 1 hour. Plates were rinsed three times with PBS 0.04% Tween-20 and incubated with 3-fold dilutions of the sampled serum starting at a 1/250 dilution during 2 hours. Plates were rinsed and further incubated with a 1/2000 dilution of an anti-mouse IgG1 (Jackson ImmunoResearch) Fc-specific conjugated with horseradish peroxidase in PBS 0.04% Tween-20 for 1 hour. After a final rinse, 3,3′,5,5′-Tetramethylbenzidine Liquid Substrate (Sigma) was used to develop the assay and the reaction was stopped after one minute using 1M HCl. Optical density was measured at 450 nm using a Modulus^TM^ II microplate reader (Turner BioSystems). Titers were calculated by logarithmic interpolation of the dilutions with readings immediately above and immediately below an OD450 of 0.2 as detailed by Ersching J et al.^25^

#### 2.4) Ageing cohorts

Ageing cohorts of *Ngfr* KO and *Bcl2 Tg* mice were established to study health issues in these models at the long term. Animals were sacrificed when reaching the humane endpoint and samples were taken. In the 35-week-cohort, animals reaching the humane endpoint before that age, were removed from the study.

#### 2.5) ANA titration assays

The detection of IgG ANAs in the serum of *Ngfr* KO and *Bcl2 Tg* mice was performed by IF in Hep-2 coated slides (Kallestad™ Bio-Rad) following manufacturer instructions. Serums were diluted at 1/50, 1/250, 1/500 or 1/1500 and a 20μl drop was applied to the assigned antigen wells together with the controls and were incubated in a moist chamber for 30 minutes at RT. Wells were rinsed and washed with PBS for 10 minutes. 20μl of conjugated anti-mouse IgG AF488 (Molecular Probes) diluted 1/200 in PBS were added to each well. Secondary antibodies were incubated and washed as indicated before for the sera and mounted for analysis. X20 images were acquired in a Nikon Eclipse Ni-E microscope (Nikon) and NIS-Elements software (Nikon).

ANAs titer was expressed as the higher dilution showing nuclear positivity for IgG and statistics were assessed using Mann-Whitney test.

### 3) Immunohistological approaches

#### 3.1) Histology and immunohistochemistry

For immunohistochemistry, tissues were sampled and fixed ON in PBS 4% formalin and stored in 50% ethanol until embedded in paraffin blocks. 1-μm thick sections were obtained from the blocks, mounted in Superfrost ® Plus microscope slides (Thermo Fisher Scientific) and dried ON. For different histological and immunohistochemical methods, slides were deparaffinized in xylene and re-hydrated through a series of graded ethanol until water.

Tissue sections were stained with H&E, or prepared for IHC in an automated immunostaining platform Ventana Discovery XT (Roche), Autostainer Link, (Dako, Agilent) or Dako Omnis (Dako, Agilent). Antigen retrieval was first performed with high pH buffer CC1m (Roche), or Low pH (Dako) depending on the primary antibody. Endogenous peroxidases were blocked using 3% hydrogen peroxide. Slides were then incubated with the appropriate primary antibody, as detailed in (**Supplementary table 6**). Afterwards, sections were incubated with the corresponding secondary antibodies (anti-mouse or anti-rat when needed) and visualization systems OmniMap anti-Rabbit, (Ventana, Roche) or Novolink Polymer, (Leica) when needed conjugated with HRP.

Immunohistochemical reaction was developed using either 3, 30-diaminobenzidine tetrahydrochloride (DAB) Chromomap DAB, (Ventana, Roche) or DAB solution (Dako) or Purple kit (Ventana, Roche) as chromogens. Carazzi’s hematoxylin was used to counterstain the nuclei. Finally, the slides were dehydrated, cleared and mounted with a permanent mounting medium (Sakura) for microscopic evaluation. Positive control tissue sections known to express the target antigen were included in each staining run.

#### 3.2) Pathological analysis of kidney damage in mice

Disease development in the *Bcl2 Tg* model was assessed by histological analysis in collaboration with the pathologist E. Caleiras (CNIO) in a blinded study.

Autoimmune glomerulonephritis was defined by the presence of hypercellular and segmented glomeruli with occluded capillaries in many cases containing eosinophilic deposits and generally accompanied by Bowman’s capsule epithelium hyperproliferation. The percentage of glomeruli affected by these last two alterations were quantified as markers of severe kidney damage in 4 X20 pictures obtained from the cortical area of each mouse in H&E stains.

### 4) Bioinformatic approaches

#### 4.1) RNAseq analysis

To perform gene expression analysis in FDCs, PO LNs from of WT and *Ngfr* KO mice were pooled to prepare homogeneous cell suspension and up to 300 FDCs were isolated by FACS and directly collected into single cell lysis buffer (TakaraBio), with 40 U/µL recombinant RNase Inhibitor (TakaraBio).

cDNA was amplified from the collected cells using the SMART-Seq v4 Ultra Low Input RNA Kit (Clontech-TakaraBio). 1ng of amplified cDNA was used to generate barcoded libraries with the Nextera XT DNA library preparation kit (Illumina). The size of the libraries was checked using the Agilent 2100 Bioanalyzer High Sensitivity DNA chip and their [EMG1] concentration was determined using the Qubit^®^ fluorometer (Thermo Fisher Scientific).

Libraries were sequenced on HiSeq 2500 and processed with RTA v1.18.66.3. FastQ files for each sample were obtained using bcl2fastq v2.20.0.422 software (Illumina). Sequencing reads were aligned to the mouse reference transcriptome (mm10 v90) and quantified with RSem v1.3.1 (31). Raw counts were normalized with TPM (Transcripts per million) and TMM (Trimmed mean of M-values) methods, transformed into log2 expression (log2(rawCount+1)) and compared to calculate fold-change and corrected p-value (FDR) using a Benjamini and Hochberg procedure. Only those genes expressed with at least 1 count in a number of samples equal to the number of replicate samples of the condition with less replicates were taken into account.

RNA-seq data generated have been deposited in the Gene Expression Omnibus under the accession number GSE236511.

The volcano plot in **Figure 4B** was generated using EnhancedVolcano R package.

For the heatmap in **Supplementary figure 4D,** the 10 top up-regulated genes for each population of stromal cells were computed using the FindAllMarkers function from the Seurat R package and used to represent RNAseq expression data from *Ngfr* WT and KO samples. Expression data were scaled for each gene before representation.

#### 4.2) Comparisons with scRNAseq datasets

Gene signature scoring was performed as previously described by Mourcin F et al.^46^ Briefly, scRNAseq data from mouse lymph node stromal cells were downloaded from the NCBI GEO database (GEO: GSE112903) and reanalyzed according to original methods in Rodda LB et al.^31^ or provided directly by the authors Pikor NB et al.^26^ Gene signatures were obtained from *Ngfr* WT and KO mice based on an exclusion criteria of a false discovery rate (FDR)<0.1 and a log2 fold change (log2FC) < or > than −1 or 1. Signature scores were computed using the AddModuleScore function from the Seurat R package (v3.1.5) with R software (v4.1.2). This function calculates for each individual cell the average expression of each gene signature, subtracted by the aggregated expression of control gene sets. All the analyzed genes were binned into 25 bins based on their averaged expression, and for each gene of the gene signature, 100 control genes were randomly selected from the same bin as the gene.

UMAPs, violin plots and heatmaps were generated using Seurat, Vioplot (v0.3.0) and pheatmap (v1.0.12) packages. Statistics in violin plots were calculated using Student’s t tests.

For **Figure 3E**, signatures for immunized and non-immunized conditions within the FDC1 cluster (Pikor et al., 2020) were computed with the FindMarkers function from the Seurat R package using Wilcoxon test (with the following parameters: avg_log2FC>0.25 or <-0.25 and p_val_adj<0.05) and used to compare with *Ngfr* WT and KO signatures.

#### 4.3) GSEA

GSEA Preranked was used to perform GSEA^88^ of Reactome pathway or custom-made databases on the pre-ranked gene list of the significantly up-regulated or down-regulated genes (FDR<0.1 and FC>1 or <-1) from the *Ngfr* WT or KO FDCs RNAseq described above. Settings were established for 1,000 gene set permutations. Only those gene sets with significant enrichment levels (FDR q value < 0.25) and more than 20 genes were finally considered.

### 5) Statistical analysis

Most graphs and statistical analyses were obtained using GraphPad Prism software version 9.4.0 (Dotmatics) and data are presented as mean ± standard deviation (SD) unless otherwise indicated. All data sets were first tested for normality using the Anderson-Darling, D’Agostino-Pearson omnibus, Shapiro-Wilk and/or Kolmogorov-Smirnov tests. Specific test used for each figure as well as biological replicates are stated in the figure legends. Survival was represented as Kaplan-Meier curves and statistics were performed using Log-rank (Mantel-Cox) test.

P-values are indicated in each figure for statistically significant comparisons (p<0.05).

## Abbreviations

ANAs: Anti-nuclear antibodies
BCL2: B cell lymphoma 2 protein
BDNF: Brain derived neurotrophic factor
BECs: Blood endothelial cells
BM: Bone marrow
BR: Brachial
DEGs: Differentially expressed genes
DNCs: Double negative cells
DZ: Dark zone
FDCs: Follicular dendritic cells
FRCs: Fibroreticular cells
GC: Germinal center
KLH: Keyhole limpet hemocyanin
LECs: Lymphatic endothelial cells
LINGO: Leucine rich repeat and Ig domain containing
LN: Lymph node
LSCs: Lymphoid stromal cells
LZ: Light zone
MAdCAM-1: Mucosal vascular addressin cell adhesion molecule-1
MRCs: Marginal reticular cells
NGF: Nerve growth factor
NGFR: Nerve growth factor receptor
NP: 4-hydroxy-3-nitrophenyl-acetyl
OS: Overall survival
PO: Popliteal
SLE: Systemic lupus erythematosus
SLOs: Secondary lymphoid organs
SRBCs: Sheep red blood cells
TfH: T follicular helper
Trk: Tropomyosin receptor kinase
VCAM-1: Vascular cell adhesion molecule-1
WT: Wild type

## Acknowledgments

We apologize to those authors whose work could not be cited due to size limitations. We thank to Dr. Miguel Angel Piris, Dr. Miguel Gallardo for their support in the project and all the members of the Microenvironment and Metastasis laboratory (CNIO) for their constant help and guidance. We also thank Dr. Veronica Matía for her help with mice experiments and Dr. Maria José Jimenez Santos for her help with bioinformatic analysis.

Dr. Peinado laboratory is funded by Project PID2020-118558RB-I00, PDC2021-121102-I00 funded by MCIN/AEI/10.13039/501100011033 and the European Union “NextGeneration EU”/PRTR, AECC (PRYCO223002PEIN), European Union through the PANCAID project, Horizon EU Programme (Grant agreement 101096309), European Union’s Horizon 2020 research and innovation programme “proEVLifeCycle” under the Marie Skłodowska-Curie grant agreement No 860303. We are also grateful for the support of La Caixa Foundation (fellowship ID100010434, LCF/BQ/ES17/11600007) and EMBO scientific exchange grant (Number 9156) awarded to A. H-B. The CNIO, certified as Severo Ochoa Excellence Centre, is supported by the Spanish Government through the Instituto de Salud Carlos III (ISCIII).

---

**Supplementary figure 1:**
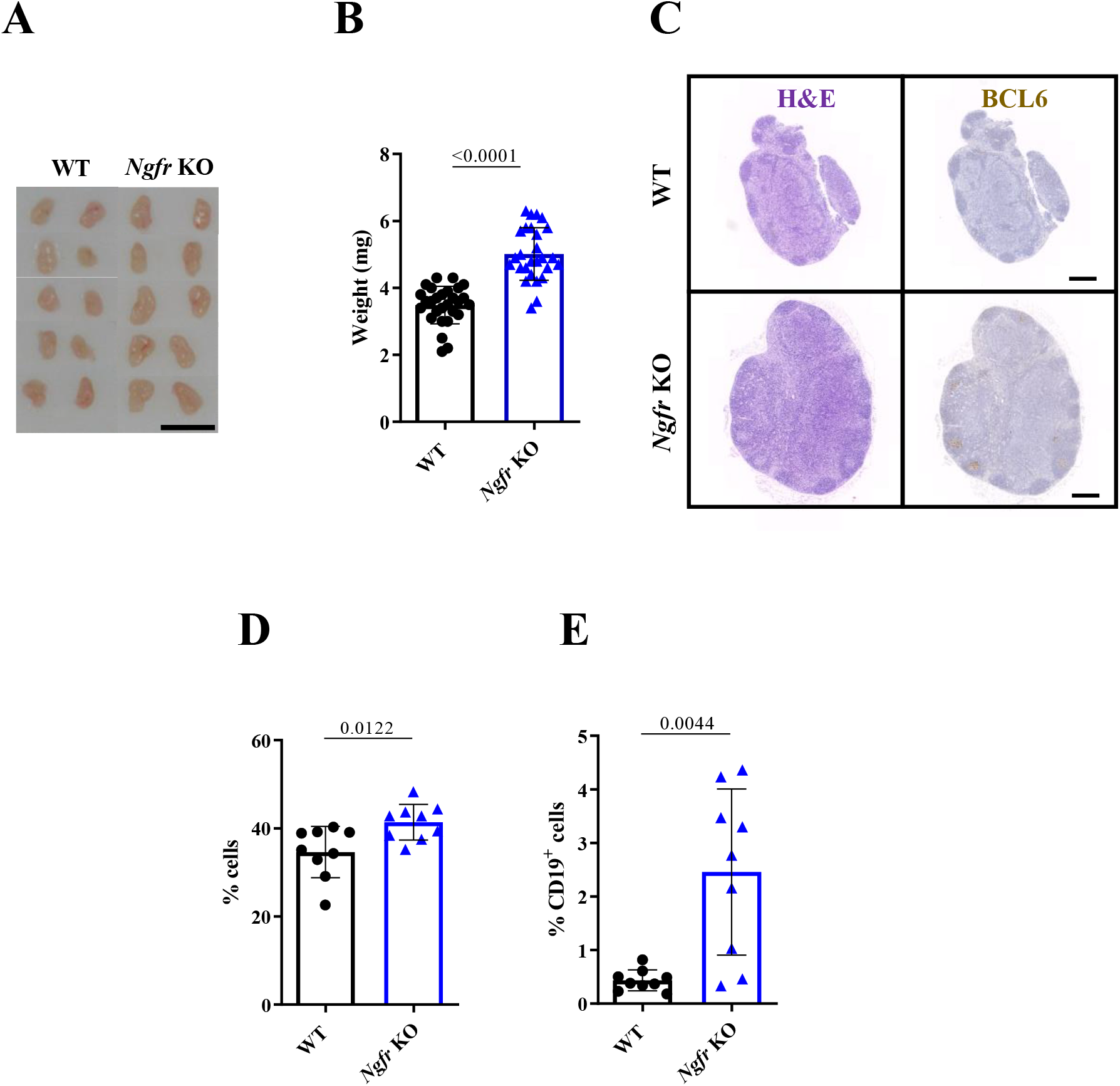
GC formation in BR LNs. **A**) Representative image of BR LN size WT and *Ngfr* KO mice. Scale bar 5mm. **B**) BR LN weight in WT and *Ngfr* KO mice. n=28 from 3 independent experiments. **C**) Representative H&E and immunohistochemistry of BCL6 in BR LNs of WT (upper panels) and *Ngfr* KO (lower panels). Scale bar, 400µm. **D**) BR LN CD19^+^ cells in WT and *Ngfr* KO mice. n=9 from 2 independent experiments. **E**) PO LN GC cells in WT and *Ngfr* KO mice n=9 from 2 independent experiments. Graphs show mean and error bars show SD. For **B**, **D** and **E** P-value by two-tailed unpaired T test with Welch’s correction.

**Supplementary figure 2:**
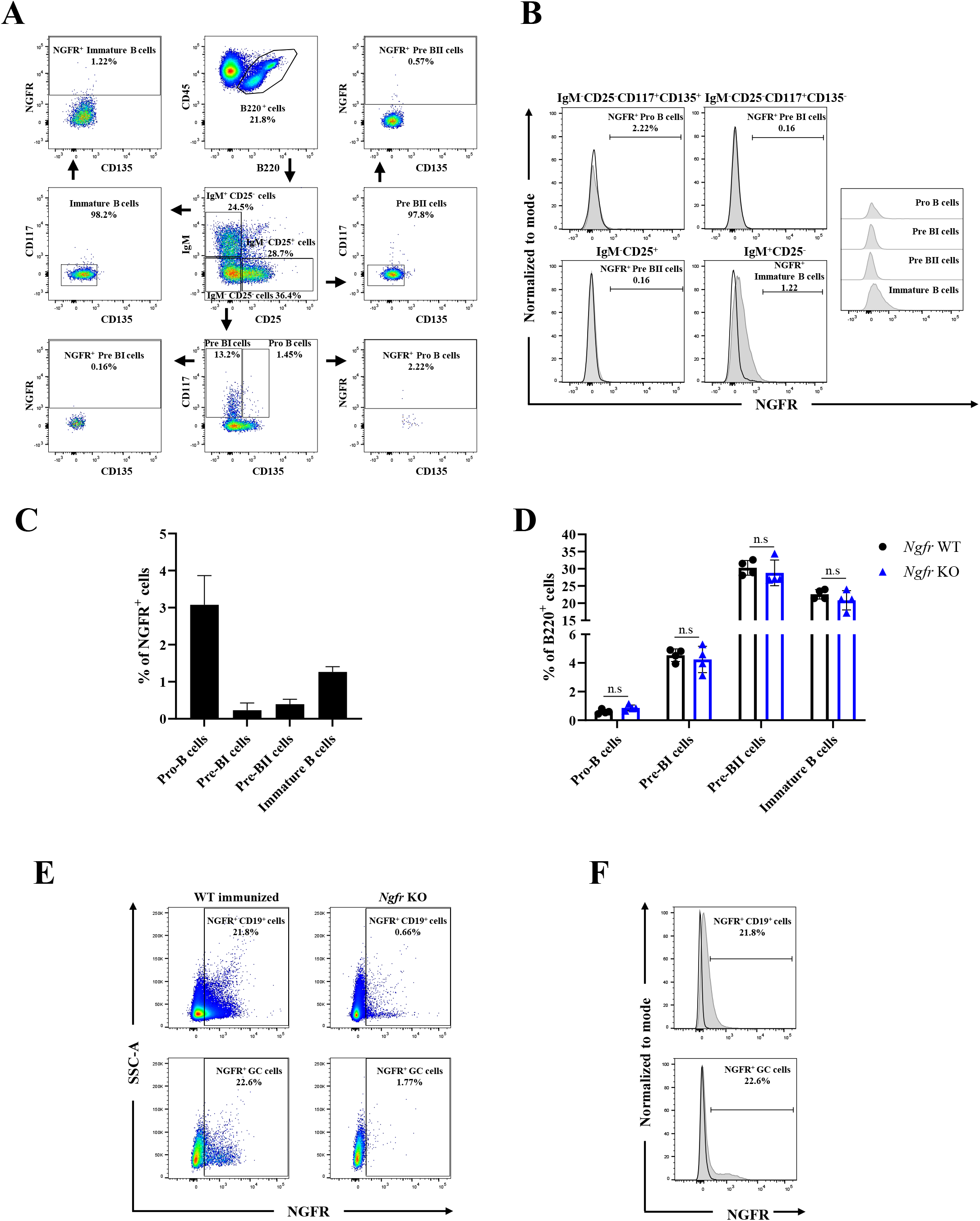
Analysis of NGFR expression in immune cells. **A**) Gating strategy to study NGFR expression in B cell precursors in the BM. **B**) Histograms comparing NGFR mean fluorescence in BM B cell precursors of WT mice gated in **A**. Black lines represent NGFR fluorescence in *Ngfr* KO BM cells used as negative CTL. **C**) Quantification of the percentage of NGFR^+^ cells in BM B cell precusors. **D**) Frequencies of the different populations of BM B cell precursors in WT and *Ngfr* KO mice. n=4 from 1 experiment. **E**) Representative plots of NGFR expression in the CD19^+^ and the CD19^+^CD95^+^CD38^lo^ GC B cells subsets in WT and *Ngfr* KO LNs by flow cytometry. **F**) Histograms comparing NGFR mean fluorescence in the populations gated in **E**. Black lines represent NGFR fluorescence in *Ngfr* KO mice used as negative CTL. Graphs show mean and error bars show SD. For **D** P-value by two tailed Mann-Whitney test.

**Supplementary figure 3:**
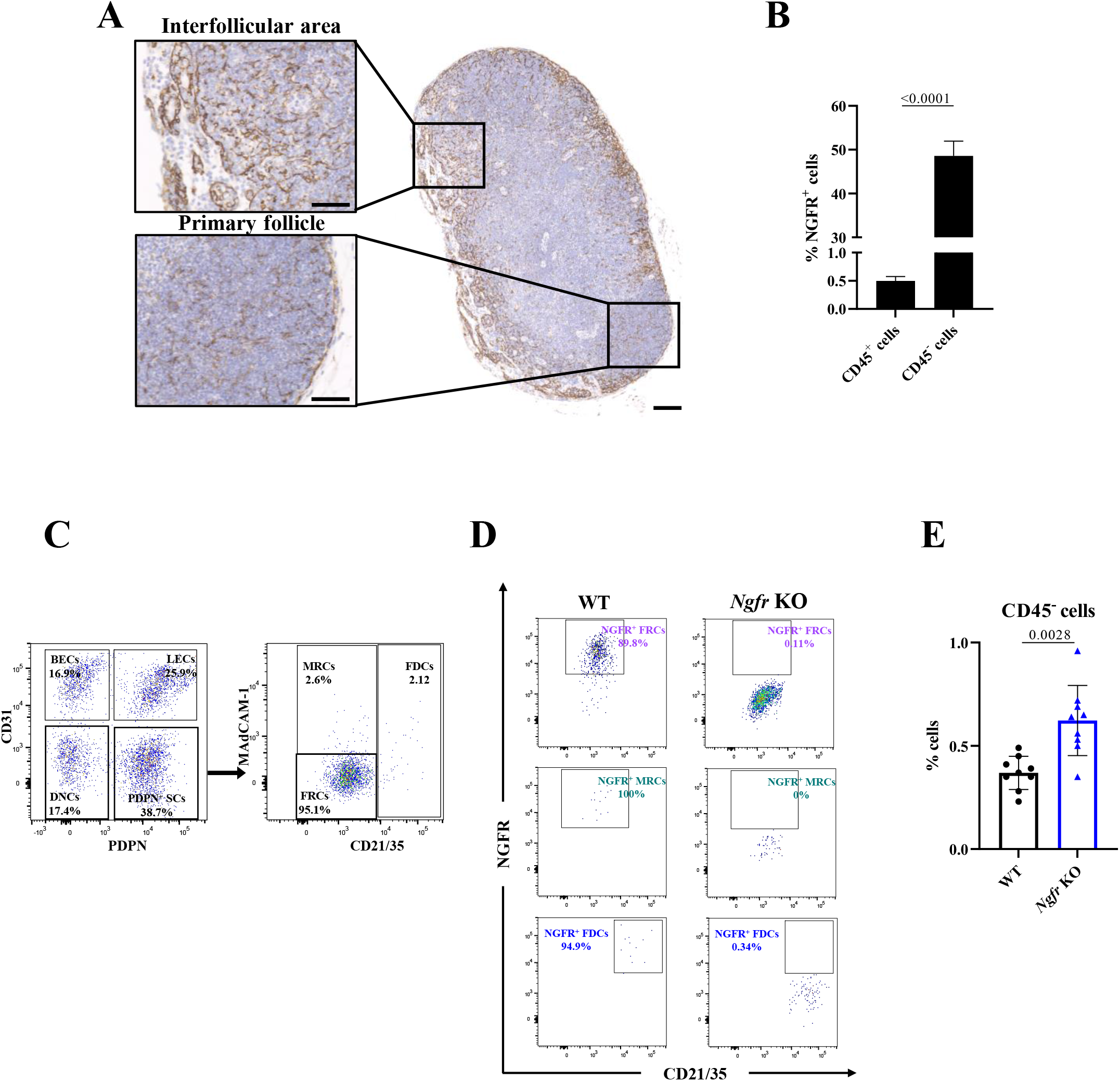
Analysis of NGFR expression in LN stromal cells. **A**) Representative IHC of NGFR in the PO LN showing the whole architecture of the organ (scale bar, 100µm) and magnification of different substructures (scale bar, 50µm). **B**) Quantification by flow cytometry of the percentage of NGFR^+^ cells in immune (CD45^+^) and stromal (CD45^−^) cell populations in PO LNs n=9 from 2 independent experiments **C**) Gating strategy of CD45^−^ LN stromal populations. **D**) Representative plots of NGFR expression in the PDPN^+^ SCs subsets in WT and *Ngfr* KO LNs. **E**) Percentage of CD45^+^ and CD45^−^ cell populations expressing NGFR in *Ngfr* WT PO LNs. n=9 from 2 independent experiments. Graphs show mean and error bars show SD. For **B** and **E** P-value by two-tailed unpaired T test with Welch’s correction.

**Supplementary figure 4:**
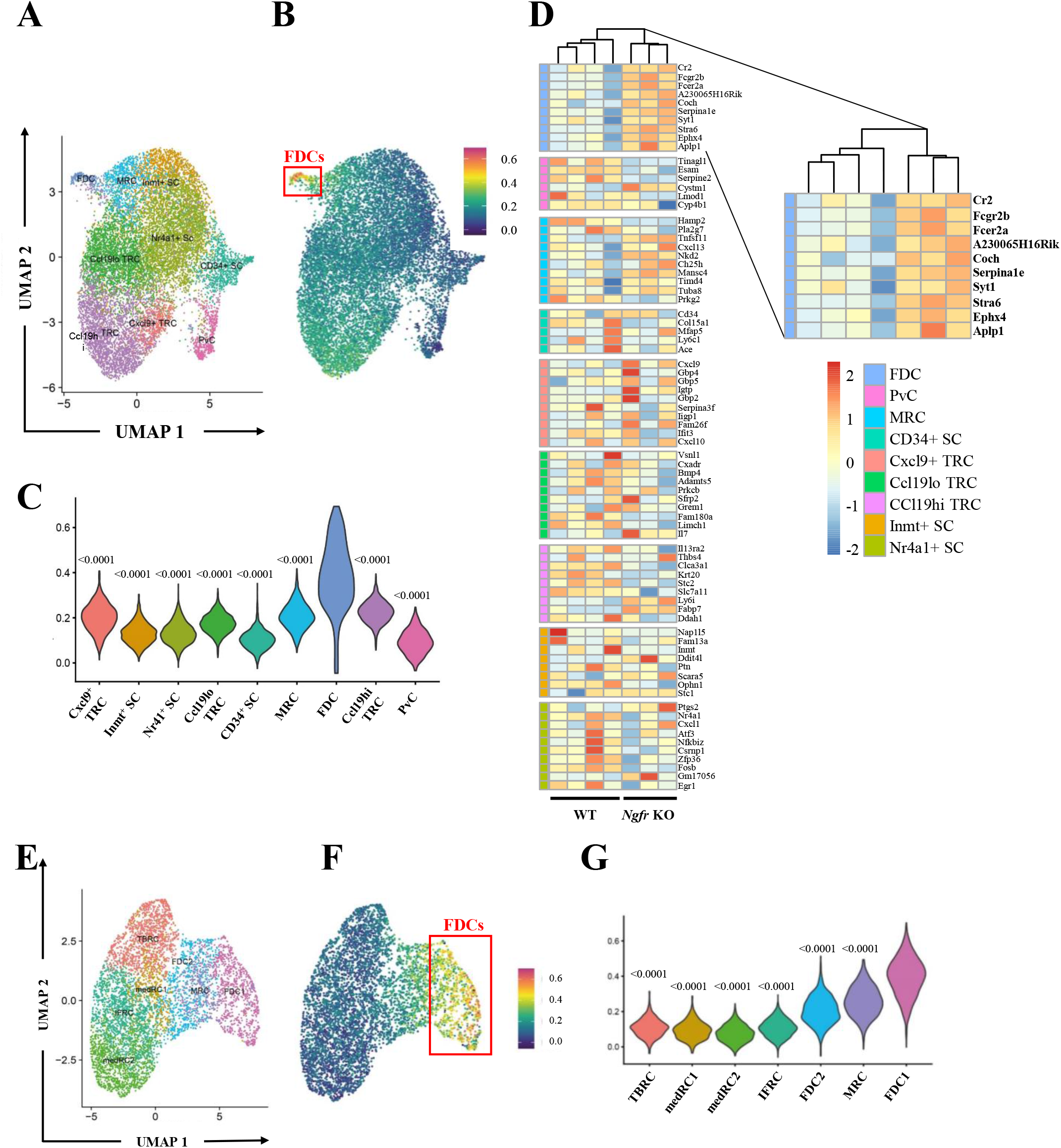
Analysis of FDC activation signatures in scRNAseq datasets. **A**) UMAP plot showing the distribution of the different subsets of stromal cells identified by scRNAseq in Rodda LB et al. **B**) Heatmap showing the comparison of the up-regulated gene signature obtained from the *Ngfr* KO mice vs WT FDCs with the clusters showed in **A**. **C**) Violin plots showing the percentage of identity of the up-regulated signature with the different cell subsets identified in **A**. Statistics refers to the comparison of each population with the FDC subset. **D**) Heatmap showing the differential expression of the RNAseq analysis of *Ngfr* KO vs WT FDCs for the top 10 genes defining each subset found by Rodda LB et al. and magnification of the FDC cluster. The color code identifies each population accordingly to the UMAP distribution in **A**. **E**) UMAP plot showing the distribution of the different subsets of stromal cells identified by scRNAseq in Pikor NB et al. **F**) Heatmap showing the comparison of the up-regulated gene signature obtained from the *Ngfr* KO mice vs WT FDCs with the clusters showed in **E**. **G**) Violin plots showing the percentage of identity of the up-regulated signature with the different cell subsets identified in **E**. Statistics refers to the comparison of each population with the FDC1 subset. For **C** and **G** P-value by two-tailed unpaired T test with Welch’s correction.

**Supplementary figure 5:**
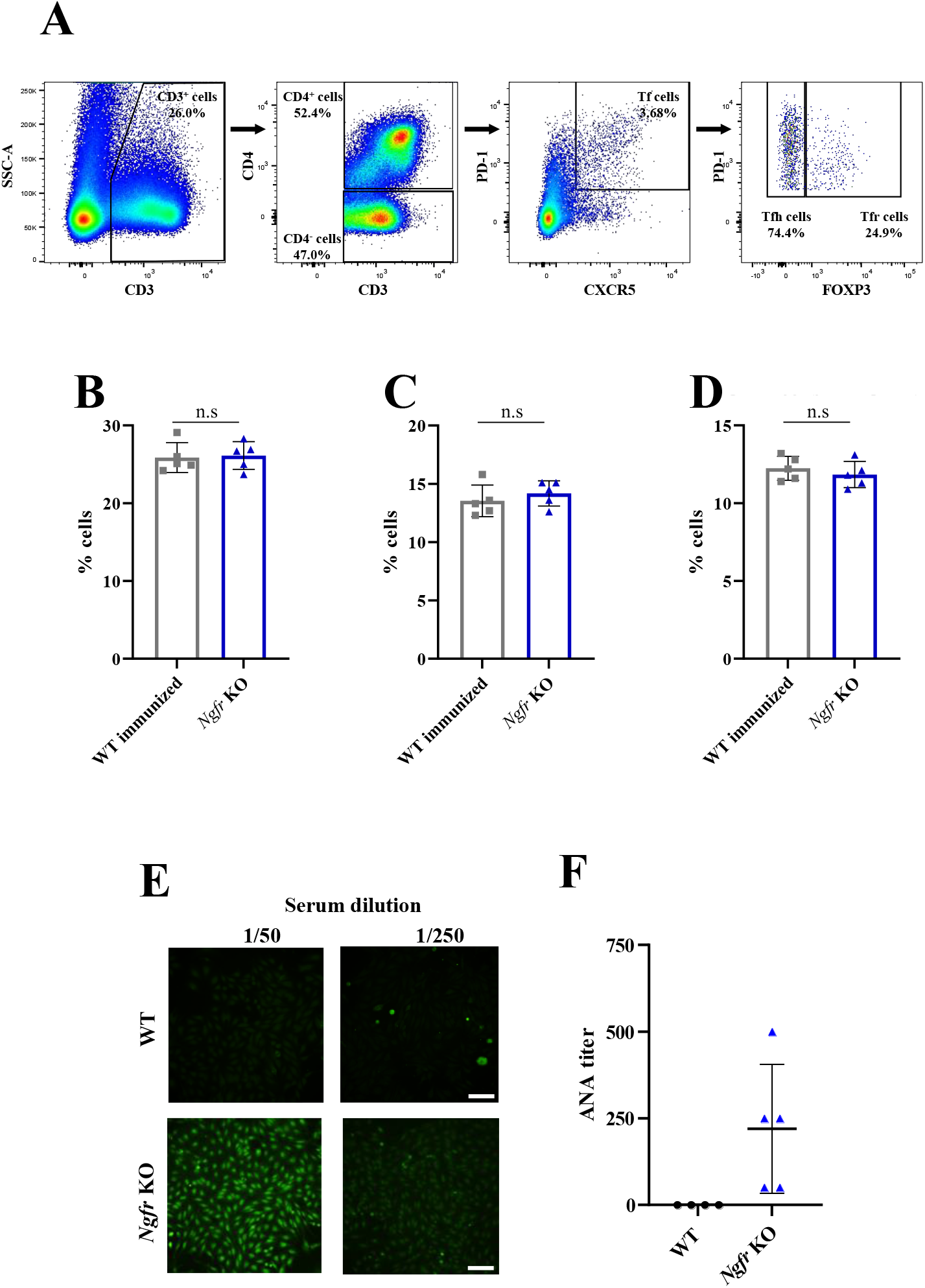
Characterization of T follicular cells and ANA production in *Ngfr* KO mice. **A**) Gating strategy used in flow cytometry analysis of LN T cells including T follicular (Tf) cells. **B)** Quantification by flow cytometry of total T cells (CD3^+^) in WT immunized and *Ngfr* KO PO LNs. **C**) Quantification by flow cytometry of CD4^+^ T cells in WT immunized and *Ngfr* KO PO LNs. **D**) Quantification by flow cytometry of CD4^−^ T cells in WT immunized and *Ngfr* KO PO LNs. **E**) Representative images from IF of Hep-2 cells slides incubated with serums of 10-week-old WT and *Ngfr* KO mice at the indicated dilutions. Scale bar, 100µm. **F**) Semi-quantitative measurement of ANAs titer in WT and *Ngfr* KO mice serums. n=4 WT and n=5 *Ngfr* KO mice. For **B**, **C** and **D** n=5 from 1 experiment. Graphs show mean and error bars show SD. For **B**, **C** and **D** P-value by two-tailed unpaired T test with Welch’s correction. For **F** Statistics are not show since all values in WT serums were 0.

**Supplementary figure 6:**
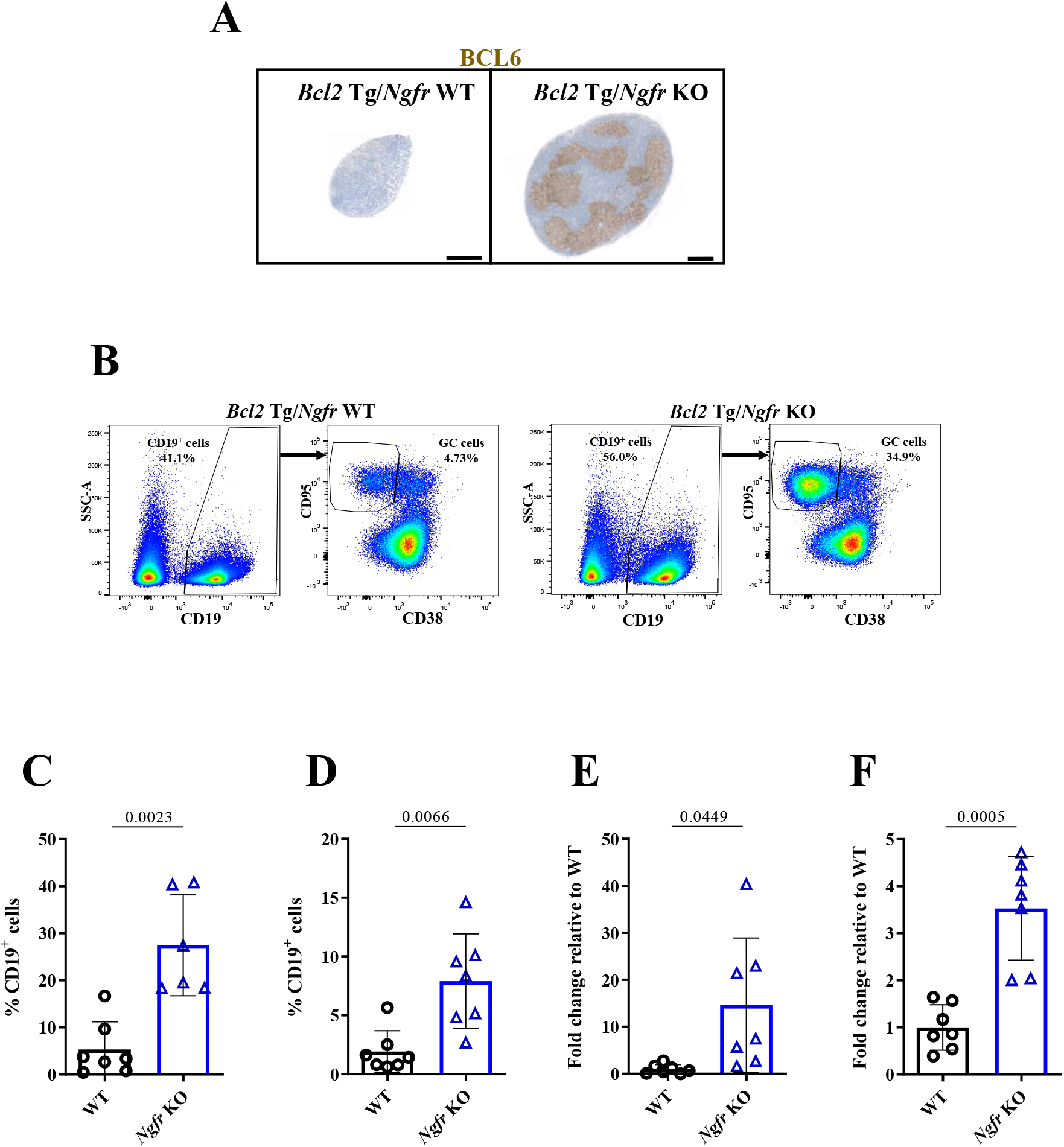
Characterization of GC formation in the *Ngfr* KO/ Bcl2 Tg model. **A**) Representative images of BCL6 IHC in PO LN from 10-week-old *Bcl2* Tg crossed with *Ngfr* WT and KO. Scale bar, 400µm. **B**) Representative examples of the gating strategy of CD19^+^ and GC cells in PO LN from *Bcl2* Tg/*Ngfr* WT (left panels) and *Bcl2* Tg/*Ngfr* KO (right panels) mice. **C)** Quantification by flow cytometry of PO LN GC cells in *Bcl2* Tg/*Ngfr* WT and KO mice. n=7 for *Bcl2* Tg/*Ngfr* WT and n=6 for *Bcl2* Tg/*Ngfr* KO from 2 independent experiments. **D**) Quantification by flow cytometry of BR LN GC cells in *Bcl2* Tg/*Ngfr* WT and KO mice. n=7 for *Bcl2* Tg/*Ngfr* WT and *Bcl2* Tg/*Ngfr* KO from 2 independent experiments. **E**) Quantification by flow cytometry of PO LN FDCs in *Bcl2* Tg/*Ngfr* WT and KO mice. Frequency of total cells is relative to the *Bcl2* Tg/*Ngfr* WT values and expressed in fold change. n=6 from 2 independent experiments. **F**) Quantification by flow cytometry of BR LN FDCs in *Bcl2* Tg/*Ngfr* WT and KO mice. Frequency of total cells is relative to the *Bcl2* Tg/*Ngfr* WT values and expressed in fold change. n=7 from 2 independent experiments. Graphs show mean and error bars show SD. For **C**, **D**, **E** and **F** P-value by two-tailed unpaired T test with Welch’s correction.

